# Accurate, high-coverage assignment of *in vivo* protein kinases to phosphosites from *in vitro* phosphoproteomic specificity data

**DOI:** 10.1101/2021.08.31.458376

**Authors:** Brandon M. Invergo

## Abstract

Phosphoproteomic experiments routinely observe thousands of phosphorylation sites. To understand the intracellular signaling processes that generated this data, one or more causal protein kinases must be assigned to each phosphosite. However, limited knowledge of kinase specificity typically restricts assignments to a small subset of a kinome. Starting from a statistical model of a high-throughput, *in vitro* kinase-substrate assay, I have developed an approach to high-coverage, multi-label kinase-substrate assignment called IV-KAPhE (“*In vivo*-Kinase Assignment for Phosphorylation Evidence”). Tested on human data, IV-KAPhE outperforms other methods of similar scope. Such computational methods generally predict a densely connected kinase-substrate network, with most sites targeted by multiple kinases, pointing either to unaccounted-for biochemical constraints or significant cross-talk and signaling redundancy. I show that such predictions can potentially identify biased kinase-site misannotations within families of closely related kinase isoforms and they provide a robust basis for kinase activity analysis.

## Introduction

Protein phosphorylation is the most common form of post-translational modification and it plays a central role in intracellular signaling. Diverse protein kinases catalyze the binding of a phosphate group to a substrate acceptor residue, typically serines (S), threonines (T) or tyrosines (Y) in eukaryotes. The active sites of kinases’ enzymatic domains exhibit phosphoacceptor-residue specificity, which can be broadly classified in eukaryotes as serine/threonine (S/T)-specific, tyrosine (Y)-specific, or so-called “dual-specificity” kinases. Kinase substrate-specificity is further determined by the protein primary and secondary structural contexts around the phosphoacceptor residue, as well as by allosteric structural mediation of docking (Ochoa *et al*., 2018; Bradley *et al*., 2021).

The sequence contexts around known phosphorylation sites (“phosphosites”) have been widely used in computational approaches to predict new substrate sites of a protein kinase. Numerous methods have been developed to achieve this, employing, for example, scoring matrices (Yaffe *et al*., 2001; Obenauer *et al*., 2003; Miller *et al*., 2008; Jung *et al*., 2010; Safaei *et al*., 2011; Wagih *et al*., 2015; Krystkowiak *et al*., 2018), neural networks (Blom *et al*., 1999, 2004; Linding *et al*., 2007), support vector machines (Kim *et al*., 2004; Dou *et al*., 2014), sequence clustering (Zhou *et al*., 2004; Xue *et al*., 2011; Wang *et al*., 2020), kinase structure (Brinkworth *et al*., 2003), or grammatical inference (Datta and Mukhopadhyay, 2015). With the emergence of phosphoproteomics by liquid chromatography and tandem mass spectrometry (LC-MS/MS) enabling the routine detection thousands of phosphorylation sites in a single experiment (von Stechow *et al*., 2015), there is now little need to predict new phosphorylation sites *in silico*. The problem has instead changed to one of *kinase-phosphosite assignment*. Accordingly, new methods have emerged, based on models such as support vector machines (Zou *et al*., 2013; Yang *et al*., 2016), multiple kernel learning (Wang *et al*., 2017), Naïve Bayes (Ayati *et al*., 2019), networks (Wagih *et al*., 2016; Ma *et al*., 2020), and knowledge graphs (Nováček *et al*., 2020). Classical scoring matrices are often still used within these or related methods to model kinase substrate specificity (Wagih *et al*., 2016; Wang *et al*., 2017; Ma *et al*., 2020; Invergo *et al*., 2020).

Two major challenges face modern kinase-substrate assignment methods. First, dependence on literature-derived annotations for model training is subject to biases towards more commonly studied protein kinases (Invergo and Beltrao, 2018). This strongly limits and biases the kinases for which assignments can be made, whereas phosphoproteomic data requires unbiased, kinome-scale assignment. Many phosphosite-prediction methods resolve this imbalance by making predictions at the level of kinase families, however these predictions will still be biased towards the features of the well-studied family members (Invergo and Beltrao, 2018). Given the increasing evidence of distinct functional roles even among closely related kinase isoforms (see, e.g., Stambolic and Woodgett (2006); Linnerth-Petrik *et al*. (2014); Hinz and Jücker (2019); Higgins *et al*. (2021)), it is exigent that phosphorylation events be resolved at the level of individual kinases. The second challenge is that many sites can be phosphorylated by more than one kinase (Hornbeck *et al*., 2015). However, a lack of complete, “all-versus-all” kinase-site training sets has prevented assignment methods from having been constructed and evaluated in such a multi-label setting. As a result, the predictive performance for promiscuous or strongly specific kinases can dominate validation metrics calculated across all kinase assignments.

Here, I describe improvements in multi-label, kinase-phosphosite assignment over previous methods for a large subset of the human kinome. I achieve this without sophisticated machinelearning methods by focusing on the above considerations and building on the latest proteome-scale, hypothesis-free data. The result is a nested model called IV-KAPhE which is built on models of *in vitro* kinase specificity and *in vivo* functional association. First, I describe the approach used to construct the model. Then I consider the hypotheses produced by it and other computational methods about the topology of the human phosphorylation network and the utility of these methods in inferring protein kinase activity from quantitative phosphoprotoemic data.

## Results

I developed IV-KAPhE over three stages. First, I sought the best sequence-based model to represent *in vitro* kinase sequence specificity. Next, I incorporated physical-interaction and structural factors that mediate phosphorylation under the *in vitro*, context-free conditions of the training data. Finally, I nested the results of the *in vitro* model with features for predicting kinase-substrate functional associations in a predictor of *in vivo* phosphorylation (Figure 1).

**Figure 1:**
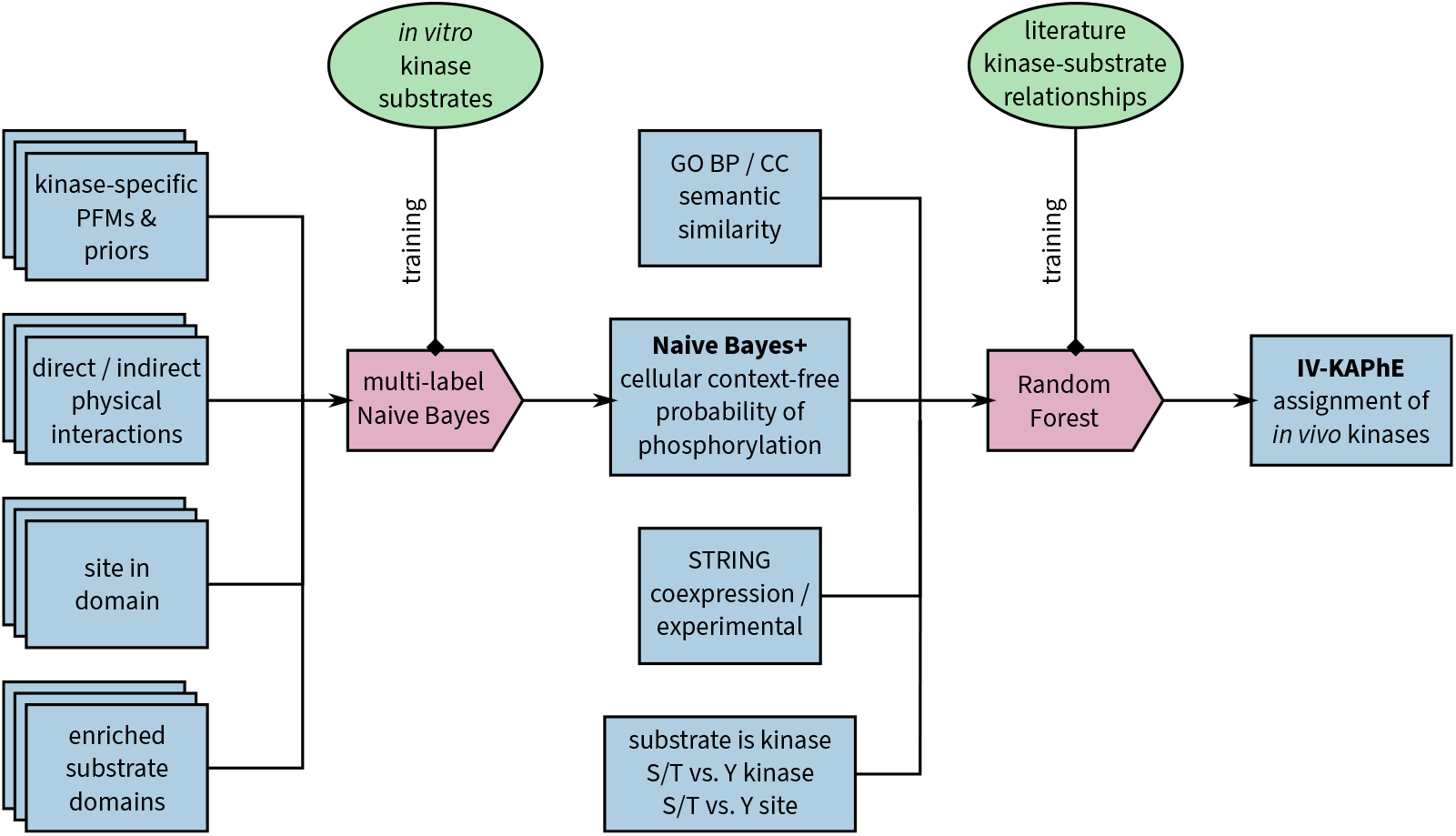
A schematic of the nested IV-KAPhE model for kinase-substrate assignment. Naïve Bayes+ consists of sub-models for each kinase, trained from kinase-specific, high-throughput *in vitro* kinase-substrate relationships. These sub-models together comprise a final, multi-label Naïve Bayes model. IV-KAPhE is a monolithic, multi-kinase Random Forest model trained from all literature-derived kinase-substrate annotations and random pairs as negative cases.

### Naïve Bayes is more appropriate than PSSM models for building a multilabel assignment method

To construct specificity models for a large fraction of the human kinome, I used results from a recent phosphoproteomic, *in vitro* assay of kinase specificity (Sugiyama *et al*., 2019). In this experiment, protein extracts from HeLa cells were first treated with a thermo-sensitive protein phosphatase and then spiked with a recombinant protein kinase. The phosphorylated extracts were digested and subjected to phosphoproteomic analysis by LC-MS/MS. This provided *in vitro* substrates for 349 protein kinases, ranging from 1 to 1672 substrates per enzyme, with 322 kinases having at least 20 substrates. As a benefit of using a single cell line, each kinase was exposed to approximately the same set of potential substrates, detectable within the limits of random sampling by shotgun proteomics.

With this data, I sought to identify the best-performing specificity model on which to build a kinase-substrate assignment method. I first considered three scoring matrix-based specificity models: the position-frequency matrix (PFM) alone, the position-specific scoring matrix (PSSM) with log-likelihood ratios of the PFM to proteomic residue frequencies, and the log-likelihood ratio PSSM backed by position-specific phosphoproteomic residue frequencies. I hypothesized that the phosphoproteome-backed PSSM would be most appropriate for the task. Next, I took advantage of the of the fact that the design of the experiment provides truenegative kinase-site assignments to reformulate PFM-based specificity as a multi-label Naïve Bayes model (Zhang *et al*., 2009). I expected that the additional prior probability information would further strengthen assignments over PSSM models.

Breaking from convention (e.g. Wagih *et al*., 2015), for performing multi-label kinase-substrate assignment I retained the central residue in scoring. This allowed me to assign S/T and Y kinases simultaneously and to handle dual-specificity kinases cleanly. To strengthen the distinction between assignment to S/T and Y kinases, I incorporated a position-weighting scheme into the models to provide greater scoring weight to highly resolved positions. I opted to use relative entropy for weighting as it provides better separation between well-defined and degenerate positions than information content does (Supplemental Figure S1). To simplify multi-label assignment, I aimed to use a single score cutoff for all kinases. Because each kinase’s PFM is unique, the scores produced by PFMs or PSSMs will have different theoretical and empirical ranges for each kinase (Supplemental Figure S2). As a result, an effective score cutoff for one kinase might be outside the theoretical range for another kinase. To overcome this, I min-max normalized the PFM and PSSM scores to be between 0 and 1 (Wagih *et al*., 2015).

I evaluated the models via 10-fold cross-validation. As the goal is to have good performance across all kinases, I chose macro-averaged precision and recall (i.e. averaged across kinases) as evaluation metrics. I avoided the Receiver Operator Characteristic (ROC) analysis commonly used in single-label prediction because the strong imbalance between positive and negative cases per kinase would de-emphasize false positives and inflate the area under the curve (Davis and Goadrich, 2006). Using these metrics, I found that raw PFMs performed poorly, while PSSM-based methods and Naïve Bayes showed overall similar performance (Figure 2a). Nevertheless, phosphoproteome-backed PSSMs out-performed proteome-backed ones, confirming that a phosphoproteome background is preferable for kinase-substrate assignment. They also performed slightly better than Naïve Bayes, against my expectations.

**Figure 2:**
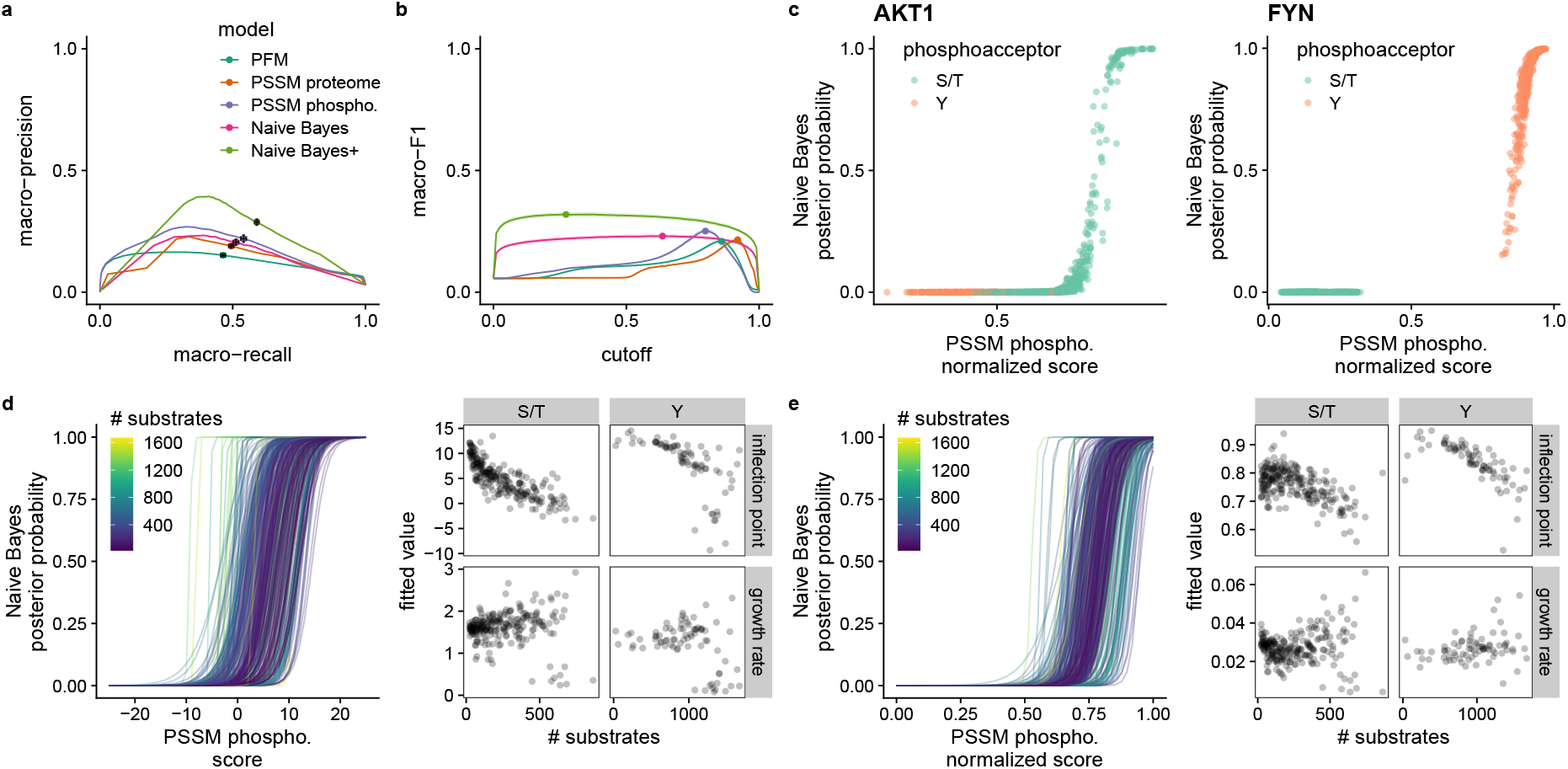
a) PSSM methods and Naïve Bayes perform similarly in cross-validation of multilabel kinase-substrate assignment via macro-averaged precision versus recall. The expanded Naïve Bayes+ model outperforms the other methods. Points indicate the scores at the cutoff that maximizes that macro-F1 score. Black error bars showing 95% confidence intervals at these points are indiscernible in most cases, indicating highly robust performance across crossvalidation folds. b) The macro-averaged F1 scores behave differently with score/probability cutoff for scoring matrix-based models versus Naïve Bayes. PSSM and PFM-based models require a strictly defined cutoff. Naïve Bayes+ again outperforms the others and retains the same flat relationship with cutoff as basic Naïve Bayes. Points indicate the maximum value. Bands indicate the 95% confidence interval. Color assignments are the same as in (a). c) Example score distributions for a S/T kinase (AKT1) and a Y kinase (FYN) from one round of cross-validation. For S/T kinases, Naïve Bayes probabilities are largely distributed close to 0.0 and 1.0 while PSSM scores take more intermediate values, notably including scores for Y sites. Y kinases show better separation for both methods. d) Left: Logistic curves relating phosphoproteome-backed PSSM scores to Naïve Bayes probabilities. Each curve represents a fitted logistic function for each kinase. The color of the curve represents the number of kinase substrates used to fit each specificity model. Right: The fitted logistic curve parameters versus number of substrates. S/T and Y kinases have negative relationships between inflection point and numbers of substrates. e) Min-max normalization of PSSM scores does not produce a stable inflection point independent of the number of substrates.

The shapes of the precision-recall curves (Figure 2a) may appear strange compared to standard dual-label curves. They can be explained first by the strong imbalance between positive and negative cases per kinase, producing very low precision at low cutoffs (i.e. at high recall). Second, at more stringent cutoffs (low recall) a growing subset of kinases is no longer assigned to any sites, in which case precision is undefined and set to 0 for each kinase so affected. Thus, the “hump” in the macro-averaged precision-recall curves (Figure 2a) represents the point at which further gains in precision from more stringent cutoffs are offset by not only lower average recall but also a reduction in effective kinome coverage.

Turning to the macro-averaged F1 score (macro-F1), which assesses the balance between precision and recall, we see different relationships between macro-F1 and cutoff (Figure 2b). Notably, macro-F1 scores of scoring-matrix methods peak at specific cut-offs before dropping precipitously, whereas Naïve Bayes gives a consistent score across much of the cutoff range. This can be explained by observing the distributions of scores or probabilities produced by the different methods (Figure 2c). While the scoring-matrix methods produce many intermediate scores (particularly for S/T kinases), Naïve Bayes mostly assigns probabilities close to 0 or 1 (Figure 2c). Moreover, the Naïve Bayes model better rejects inappropriate phosphoacceptors (e.g. Y phosphoacceptors for S/T kinases) in this way (Figure 2c). Thus, the PFM and PSSM-based models are particularly sensitive to the chosen score cutoff.

Naïve Bayes has the convenient property of a well-defined probability cutoff for assignment for all kinases (P > 0.5; see Supplemental Methods). No such definition exists for the PSSM model. Given the PSSM models’ sensitivity to cutoff selection, I sought to determine a robust selection method. As illustrated in Figure 2c, there is a sigmoidal relationship between phosphoproteome-backed PSSM scores and Naïve Bayes posterior probabilities, which could possibly be used to produce a PSSM score cutoff analogous to 0.5 posterior probability. Indeed, the phosphoproteome-backed PSSM score can be approximated by a logit transformation of the Naïve Bayes posterior probability (see Supplemental Methods). This approximation includes a dependency on the number of substrates used to fit a model, such that the 0.5-equivalent cutoff decreases with increasing number of substrates. In other words, because of varying training-set sizes, the log-likelihood PSSM score at which the foreground evidence effectively outweighs the background evidence also varies, whereas this is stabilized through normalization against total probability in Naïve Bayes.

To verify this, for each kinase I fit a logistic curve to their PSSM scores versus Naïve Bayes posterior probabilities calculated on a high-confidence set of human phosphosites (Ochoa *et al*., 2020) (Figure 2d). The fitted inflection point provides the PSSM score that is equivalent to a posterior probability of 0.5. As predicted, I observed a strong dependence between this inflection point and the number of substrates used to fit the models, which differed with kinase type. I then checked whether min-max normalization of the PSSM scores remedies the problem (Figure 2e). For S/T kinases, the inflection points were generally around 0.75, which is close to the observed macro-F1-maximizing cutoff of 0.803 (Figure 2b). However, both kinase types still had a decreasing inflection point with increasing number of substrates.

Together, these results suggest first that a raw PFM (as used in, e.g., Yang *et al*., 2016) is the weakest model. PSSMs perform best with a position-specific phosphoproteome background, but they require kinase-specific cutoffs, even after normalization, for multi-label assignment. The Naïve Bayes model offers good performance and a stable and universal cutoff. It is thus a better foundation for incorporating other features.

### Physical-interaction and structural features improve *in vitro* Naïve Bayes predictive performance

Kinases can be optimized to bind with their substrates by physical interactions outside the enzymatic kinase domain’s active site, and structural features of the substrate might further impact these interactions. These would remain relevant in the *in vitro* experimental environment and thus may be exploited to improve the model. Given the Naïve Bayes assumption of feature independence, it is trivial to incorporate other features. Thus, I constructed a second Naïve Bayes model (“Naïve Bayes+”) with added features based on: proteins physically interacting, directly or indirectly, with the kinase; proteins carrying a domain that is enriched among the kinase’s substrates or interactors; and the phosphosite being within a protein domain. These features were modeled as Bernoulli-distributed, and each showed an overall difference across the kinome in empirical probability among substrates versus non-substrates (Figure 3). As with calculating residue frequencies in substrate sequences, these empirical probability estimations may be impacted by stochastic detection of low-abundance substrates by LC-MS/MS, particularly for kinases with few substrates.

**Figure 3:**
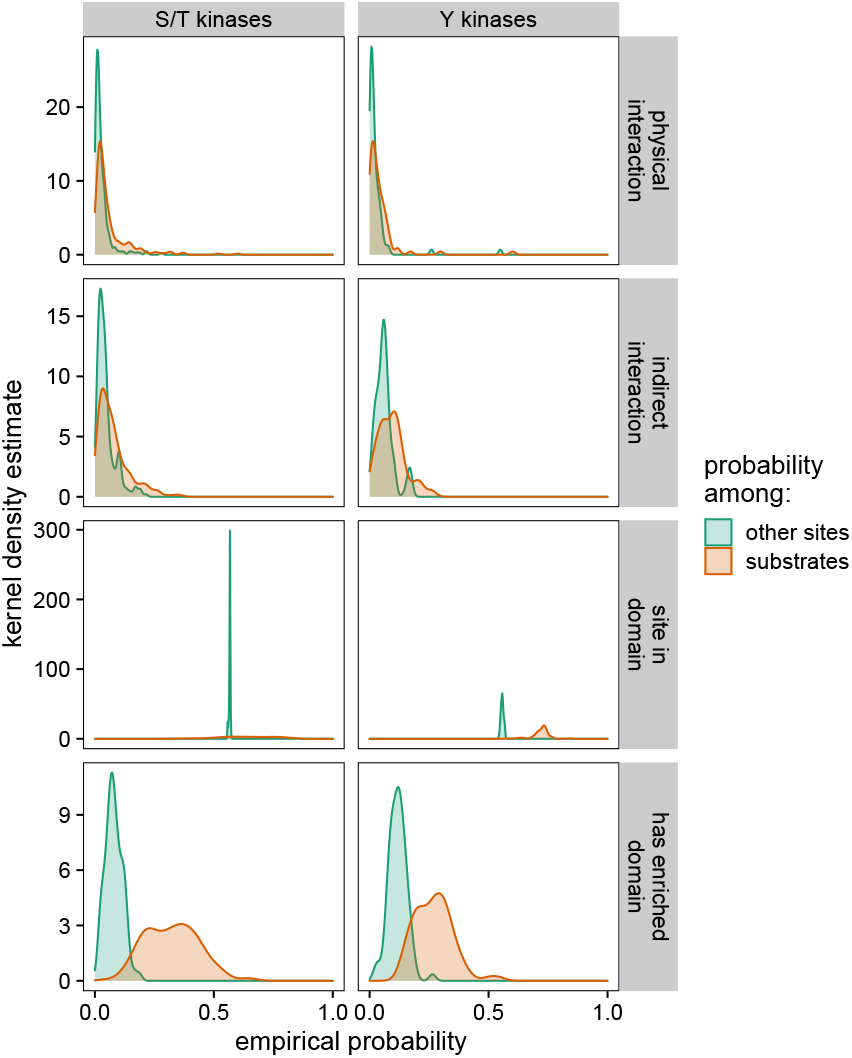
Distributions of empirical probabilities of four additional predictive features of *in vitro* kinase substrates. Distributions are represented by the kernel density estimate of the kinases’ respective Bernoulli probability parameters. Each feature trends towards higher probability in *in vitro* substrates than in other sites.

In cross-validation, the Naïve Bayes+ model produces superior macro-averaged precision and recall than the other methods (Figure 2a). Its F1 score is similarly higher, while exhibiting the same flat behavior as the sequence-only Naïve Bayes model (Figure 2b). Given this improvement in performance I carried the Naïve Bayes+ model forward for nesting into the *in vivo* model.

### IV-KAPhE produces accurate multi-label assignment of *in vivo* kinases to phosphosites

I constructed an *in vivo* kinase-substrate assignment method using the Random Forest model on five features for protein-protein functional association (Figure 1): the kinase-specific Naïve Bayes+ posterior probability for the site; the semantic similarities between the Gene Ontology (GO) “biological process” (BP) and “cellular component” (CC) annotations of kinase and substrate; and coexpression and high-throughput experimental scores between kinase and substrate from the STRING database (von Mering *et al*., 2005; Szklarczyk *et al*., 2019). Other features from STRING were discarded either by feature importance analysis (gene fusion, genome co-occurence, conserved neighborhood) or to avoid potential circularity with the training set (database imports, text-mining, combined score). I added three further features to strengthen possible discrepancies in performance (Figure 1): whether or not the substrate is itself a kinase, necessary because the functional association scores are symmetrical and can therefore produce false positives if the enzymatic roles are, in fact, reversed; whether the kinase is a S/T or Y kinase, to account for kinase-type differences in Naïve Bayes+ probability distributions; and the phosphoacceptor type (S/T or Y), for similar reasons. Because there are relatively few features and none of these require any special consideration, other machine learning models could be substituted in for *in vivo* prediction. Here, I chose Random Forest for its simplicity and for its facility in the analysis of feature importance.

For brevity, I will call this model “IV-KAPhE” (“*In Vivo*-Kinase Assignment for Phosphorylation Evidence”). For training IV-KAPhE, I took advantage of the fact that all kinase-specific substrate specificity information was used when training the underlying Naïve Bayes+ model. I thereby trained IV-KAPhE as a monolithic, non-kinase-specific model on the 7322 human *in vivo* kinase-substrate relationships annotated in the PhosphoSitePlus database (Hornbeck *et al*., 2015) and on an equally sized set of random assignments of kinases to human phosphorylation sites (Hornbeck *et al*., 2015; Ochoa *et al*., 2020) as negative cases. During training, feature importance analysis revealed the Naïve Bayes+ posterior probability, the STRING “experimental” score, and GO BP semantic similarity to be the most important (Figure 4a). The features accounting for substrate kinases, S/T versus Y kinases, and S/T versus Y sites carried low importance but their omission worsened performance, particularly for Y kinases.

**Figure 4:**
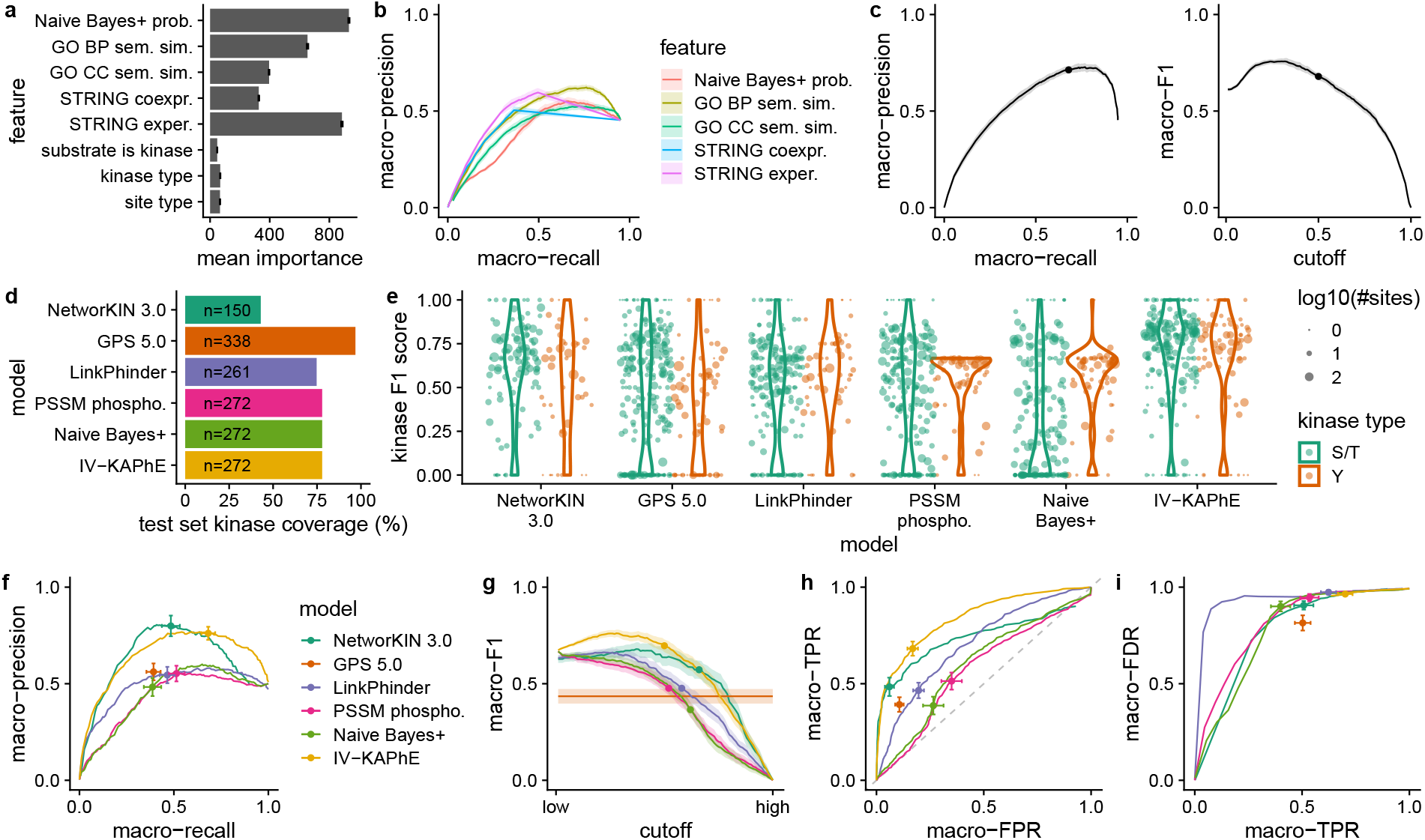
a) Naïve Bayes+ posterior probability, GO BP semantic similarity, and STRING experimental score had the greatest importance when training the Random Forest models. Error bars show standard error across cross-validation runs. b) Predictive performance of individual quantitative features, as assessed by average macro-precision and macro-recall across 10 folds of the training data, reveals GO BP semantic similarity and STRING experimental score as being the most predictive individual features. c) Cross-validation evaluated via macro-averaged precision, recall and F1 all reflect strong performance by IV-KAPhE. d) IV-KAPhE’s coverage of the external test data set is similar to LinkPhinder’s but is lower than that of GPS 5.0. e) Kinase-specific F1 scores reveal IV-KAPhE’s consistently strong performance across most kinases, with similar performance for S/T and Y kinases, compared to other methods. f) IV-KAPhE outperforms the simpler PSSM-based and Naïve Bayes+ methods as well as other previously published methods in kinase-substrate assignment of an external validation set. Points indicate the scores for simple assignments (GPS) or the scores at nominal cutoffs for quantitative predictions (cutoffs – IV-KAPhE: 0.5, PSSM: 0.75, Naïve Bayes+: 0.5, LinkPhinder: 0.672 (Nováček *et al*., 2020), NetworKIN 3.0: 1.0 (Horn *et al*., 2014)). Error bars show the 95% confidence intervals at these points. g) IV-KAPhE has a higher macro-averaged F1 score than the other methods. Points and color assignments are as in (e). Bands indicate the 95% confidence interval. h) IV-KAPhE similarly outperforms the other methods in Receiver Operating Characteristic (ROC) curve analysis for this balanced test set. Points and color assignments are as in (e). Error bars show 95% confidence intervals. i) Focusing on multi-label assignment for sites in the test set with known kinases, the macro-averaged false discovery rate (FDR; i.e. rate of novel assignments) dominates the average true positive rate (TPR). The curves are similar for most methods. At its nominal cutoff, IV-KAPhE has the highest FDR, but it is matched by the highest TPR.

While IV-KAPhE was built as a multi-label method, the training set does not represent a full, all-*versus*-all compendium of kinase-phosphosite assignments among the represented kinases and sites. This prevents full evaluation of the method in a true multi-label setting. I nevertheless evaluated it and the individual features using multi-label metrics as described above. Assessing the quantitative features’ individual predictive performances via macro-precision and macro-recall on 10 folds of the training data, GO BP semantic similarity and the STRING “experimental” score showed the highest performance (Figure 4b). Overall, the full model showed strong performance in cross-validation, reaching an average precision of 0.713 and recall of 0.679 at its nominal probability cutoff of 0.5 and outperforming the individual features (Figure 4c). The macro-F1 score at this cutoff, 0.679, was near-maximal (Figure 4c). Relaxing the cutoff slightly would improve recall, and thus F1, without significant loss of precision. To verify that the choice of machine learning model does not strongly impact performance, I repeated the cross-validation analysis, substituting a Support Vector Machine for the Random Forest model. Performance was overall quite similar (macro-precision: 0.676; macro-recall: 0.620; macro-F1: 0.626), confirming that the choice of *in vivo* model is indeed flexible.

I next evaluated IV-KAPhE’s performance on an external data set. To achieve this, I collected kinase-substrate relationships identified using the ProtMapper method from databases and textmining (Bachman *et al*., 2019). I omitted any relationships present in the PhosphoSitePlus database, from *in vivo* or *in vitro* experiments, to avoid validating on sites used in training IV-KAPhE or other methods. I then matched these 6199 previously unseen relationships with an equal number of random kinase-site relationships.

I compared IV-KAPhE to the *in vitro* Naïve Bayes+ and phosphoproteome-backed PSSM models, as well as to other previously published methods with similar or better kinome coverage (Supplemental Figure S3). To the best of my knowledge, the methods that meet that criterion are GPS 5.0 (Wang *et al*., 2020) and LinkPhinder (Nováček *et al*., 2020), both of which have greater coverage than IV-KAPhE. I also compared it to NetworKIN 3.0 (Linding *et al*., 2007; Horn *et al*., 2014), which has much lower coverage, but it shares a similar *in vitro/in vivo* hierarchical structure as IV-KAPhE. It must be stressed that none of these previously published methods were developed for multi-label assignment. Nevertheless, in order to compare their performance to IV-KAPhE, they will herein be evaluated under a multi-label paradigm. Assignments from GPS were as selected by the software’s default, “medium”-stringency behavior, because each kinase requires a different score cutoff. For LinkPhinder and NetworKIN, I evaluated performance over a range of cutoffs, with a focus on LinkPhinder’s published high-stringency cutoff of 0.672 (Nováček *et al*., 2020) and NetworKIN’s nominal likelihood-ratio cutoff of 1.0 (Horn *et al*., 2014). Each method covered a different, incomplete subset of the kinases in the test set (Figure 4d), with GPS 5.0 having the greatest coverage. NetworKIN has a significantly smaller coverage than the other methods. In order not to penalize models for coverage, each one was evaluated only on the subset of kinases that it could assign.

Like the training set, this test set is likely incomplete and cannot be fully evaluated in an all-versus-all multi-label sense. Therefore, only those relationships explicitly annotated in the test set were evaluated. I first looked at per-kinase F1 score performance, which underlies the macro-averaged metric (Figure 4e). From this view, it is clear that IV-KAPhE produces largely consistent, high F1 performance across both S/T and Y kinases compared to the other methods. NetworKIN, GPS, and LinkPhinder all exhibit highly varied performance, with GPS notably showing weaker performance for Y kinases. PSSMs and Naïve Bayes+ likewise show varied performance and weak Y-kinase performance. Note that kinases with few test-case sites tend to cluster near 0 and 1 due to lack of resolution in calculating precision and recall.

In macro-averaged precision and recall, LinkPhinder and GPS 5.0 performed only as well as the simpler, *in vitro* PSSM and Naïve Bayes+ models (Figure 4f). NetworKIN and IV-KAPhE together showed the best precision, but IV-KAPhE provided it with superior recall and averaged across a much larger portion of the kinome (Figure 4f). Returning to the macro-F1 score, we similarly see that IV-KAPhE better balances precision and recall than the other methods (Figure 4g). As this test set is balanced for each kinase between positive and negative cases, a ROC analysis is feasible (Figure 4h). Comparing the macro-averaged false-positive and true-positive rates provides further evidence that IV-KAPhE (AUC = 0.833) out-performs the other methods (AUC: LinkPhinder = 0.705; NetworKIN 3.0 = 0.673; Naive Bayes+ = 0.599; PSSM = 0.572; as a range of cutoffs were not tested for GPS 5.0, no AUC could be calculated).

I next evaluated how many novel assignments are generated by the methods when performing all-versus-all kinase-site assignment. Considering only the kinases assignable from the test set and only the test-set sites for which at least one true kinase had been assigned, I compared the macro-averaged false discovery rate (FDR) and true positive rate (TPR) across the different methods. Here “false discovery rate” is a misnomer, as we do not know whether these assignments are true or false. For all models, FDR dominates the TPR: while the fraction of known cases correctly assigned may be high, the fraction of assignments that are novel is much higher (Figure 4i). IV-KAPhE’s high FDR is nevertheless matched by a higher TPR than the other models. Thus, although IV-KAPhE also produces many novel assignments, we have a greater expectation of precision in its predictions

### Computational assignments hypothesize widespread signaling cross-talk and redundancy

The topology of the human phosphorylation network is largely unresolved and biased towards commonly studied kinases (Hornbeck *et al*., 2015; Invergo and Beltrao, 2018). One open question is to what degree sites are phosphorylated by few kinases, as illustrated in canonical signaling pathway maps, versus multiple kinases through signaling noise, redundancy, or cross-talk between pathways. By applying a near kinome-scale kinase-site assignment model to a set of phosphosites representative of the phosphoproteome, we can produce hypotheses for such questions. Accordingly, I used Naïve Bayes+ and IV-KAPhE to generate all-versus-all kinase-substrate assignments for a set of 271432 unique human phosphosites (*P* > 0.5 assignments in Supplemental Table S1; a full, unfiltered table has been archived at doi: 10.5281/zenodo.6325198), derived from the union of the entire PhosphoSitePlus human phosphosite set (Hornbeck *et al*., 2015) and a high-confidence human phosphoproteome (Ochoa *et al*., 2020). I compared these to literature-derived assignments from PhosphoSitePlus and to LinkPhinder’s assignments for human phosphosites in PhosphoSitePlus (high-stringency cutoff).

Most sites with a causal kinase annotated in PhosphoSitePlus have a single kinase annotated to them and very few have more than four annotated kinases (Figure 5a). Thus, from the literature, we would suppose that multiple kinases rarely phosphorylate the same site. However this resource contains little data for understudied kinases such as isoforms of more commonly studied ones, which tend to have similar sequence specificities (Invergo and Beltrao, 2018; Bradley and Beltrao, 2019).

**Figure 5:**
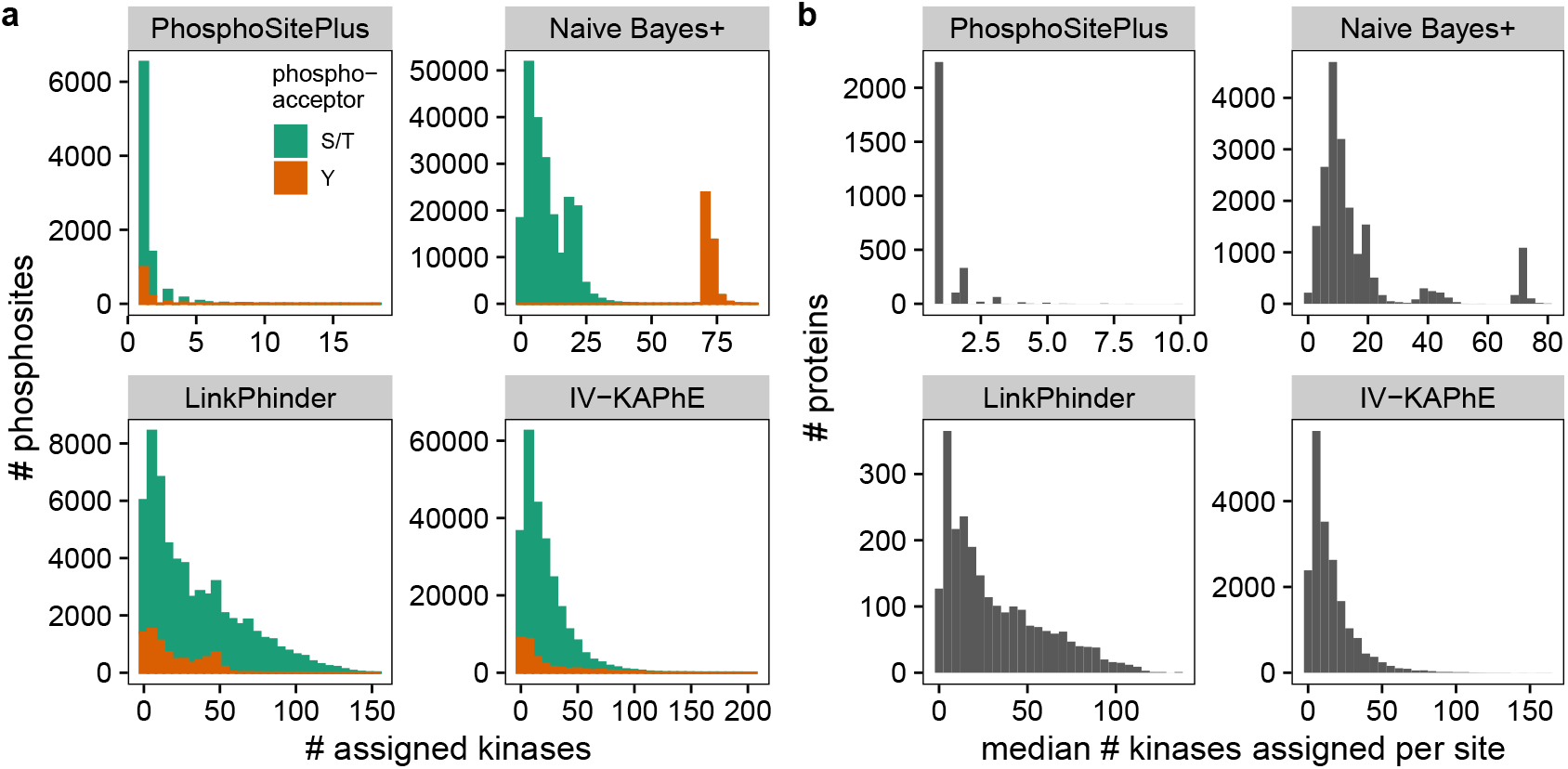
a) Histograms of the number of kinases associated with sites in the phosphoproteome reveal different views of the phosphorylation network. Literature annotations in PhosphoSitePlus suggest most sites are regulated by one or two kinases. *In vitro* Naïve Bayes+ predicts some S/T sites are “hubs” and all Y sites can be phosphorylated by most Y kinases. LinkPhinder and IV-KAPhE, in contrast, predict a long tail of hub sites. b) Histograms of the median number of kinases assigned per site for all proteins likewise show different predictions for hub proteins. Literature annotations suggest most proteins are phosphorylated by one kinase at each site. The computational methods all hypothesize multiple kinases per site, with some substrate proteins being very promiscuous at all their sites.

In contrast, Naïve Bayes+ predicts that most S/T sites are phosphorylated by multiple kinases, but generally fewer than 10 (Figure 5a). However, it also predicts that a subset of S/T sites can be phosphorylated by around 20 different kinases. Conversely, the method assigns most tyrosine kinases to each Y site, pointing to a clear technical shortcoming of the *in vitro* model: cellular context plays the major role in determining Y kinase specificity. LinkPhinder and IV-KAPhE, on the other hand, both produce long-tailed distributions, with some sites having over 100 kinases assigned to them (Figure 5a). Reassuringly, the multi-modal distribution of Naïve Bayes+ is smoothed out by the biological context incorporated into IV-KAPhE, and it assigns fewer kinases to Y sites than Naïve Bayes+.

We can similarly ask whether specific proteins act as signaling hubs between pathways, with multiple kinases phosphorylating each of their phosphosites. To answer this, I compared the median number of kinases assigned to sites for each substrate protein (Figure 5b). Again, literature-based assignments in PhosphoSitePlus suggest that most proteins are phosphorylated by a single kinase at each site. All three computational methods, on the other hand, propose the hypothesis that many proteins can be phosphorylated by multiple kinases at each of their sites (Figure 5b), i.e. that functional hubs on the protein-protein functional association network encounter many kinases, each with potentially similar, degenerate sequence specificity to the others.

One possible technical explanation of this for IV-KAPhE is that hub proteins’ strong functional association scores may override low Naïve Bayes+ probabilities via some branches in the Random Forest model. This could arise from low, false-negative Naïve Bayes+ probabilities in the training set. As a result, IV-KAPhE would produce false positives for proteins occupying central positions in the network. For sites on such proteins, a *post-hoc*, stringent filter could be applied to select only assigned kinases with high Naïve Bayes+ scores.

### IV-KAPhE predictions identify possible misannotations of kinase isoform activity

The composition of the human kinome is the result of extensive duplication events and is thus defined by families of kinases with highly similar sequence specificities (Bradley and Beltrao, 2019). Among closely related isoforms, often only one or two receive significant research attention, leading to an imbalance in kinase-substrate annotations among them (Invergo and Beltrao, 2018). Taking advantage of IV-KAPhE’s broad kinome coverage, which is less biased in composition than literature annotations, I investigated patterns of substrate assignment among closely related isoforms.

First, I considered ribosomal protein S6 kinase alpha (S6K-*α*) isoforms, among which isoforms 1 and 3 are the most commonly studied (Invergo and Beltrao, 2018). Using the full phospho-proteome assignments described in the previous section, I collected IV-KAPhE assignments of S6K-*α* isoforms for any site annotated in PhosphoSitePlus as being phosphorylated, *in vitro* or *in vivo*, by at least one of them (Supplemental Table S2; Figure 6a). The bias towards annotations for S6K-*α*-1 and −3 is plainly visible. There are a number of sites that IV-KAPhE predicts as being putative substrates of all six isoforms, pointing to multi-kinase phosphorylation, as well as some that are not well predicted for any isoform.

**Figure 6:**
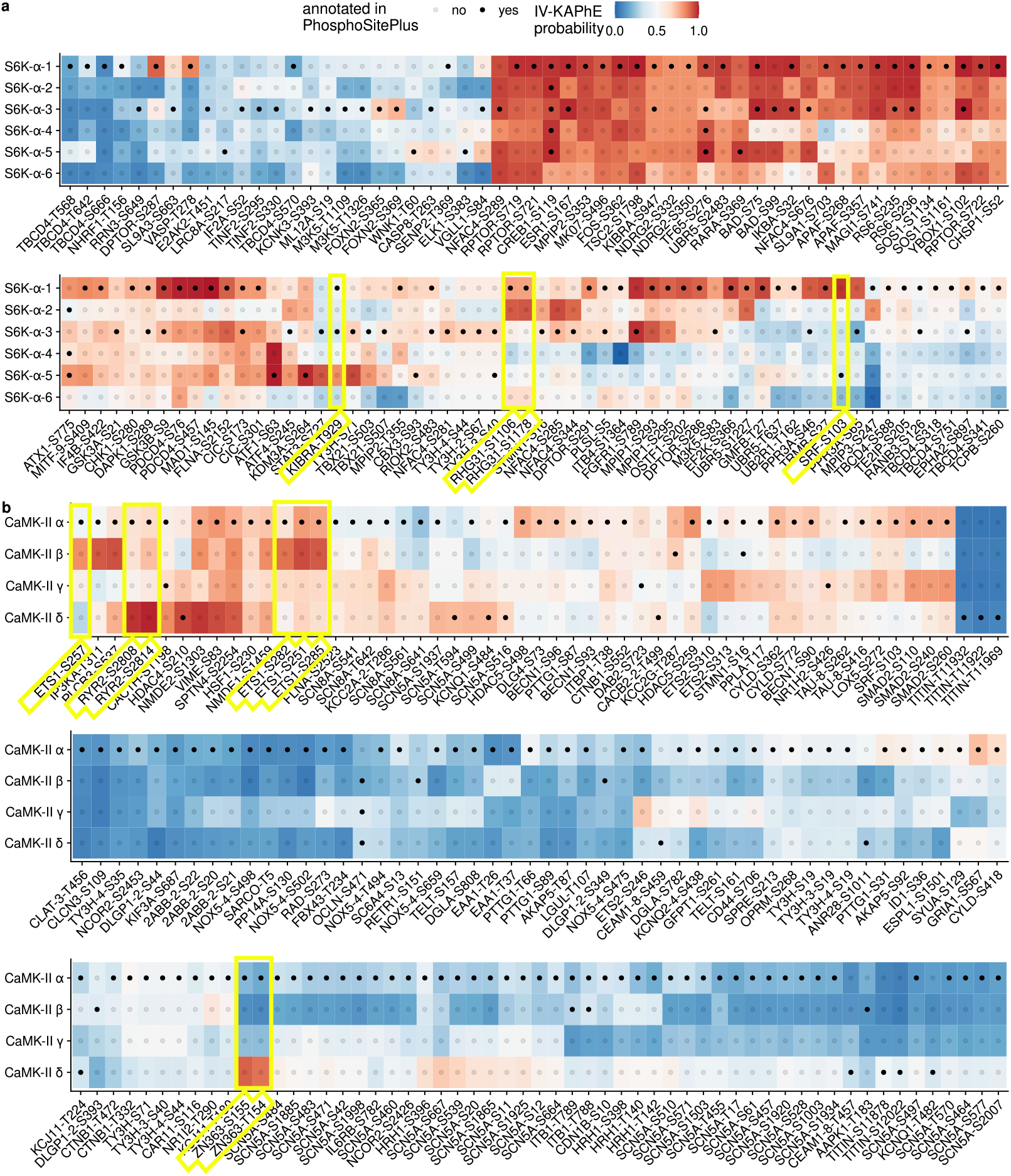
a) IV-KAPhE assignments of ribosomal protein S6 kinase alpha isoforms for all sites annotated in PhosphoSitePlus as being phosphorylated by at least one of the isoforms. Red colors indicate assignments predicted as likely by IV-KAPhE. Highlighted sites, discussed in the text, are examples that IV-KAPhE predicts are more likely to be phosphorylated by a different isoform than the one annotated. b) IV-KAPhE assignments of calcium/calmodulin-dependent protein kinase type II subunits to annotated sites, as described in (a).

More interestingly, there are sites for which an isoform has stronger probability of phosphorylating the substrate than the annotated isoform(s), supporting the argument that multi-label assignment should be carried out for kinases themselves rather than for kinase families or higher levels of classification. For example, site S103 on serum response factor (*SRF*; Uniprot: P11831 / SRF HUMAN) is annotated as a substrate of isoform 5, whereas isoform 1 has the strongest evidence (Figure 6a). This annotation was derived from an *in vitro* study, in which the isoform used was not specified (Rivera *et al*., 1993). Two sites on Rho GTPase-activating protein 31 (*ARHGAP31*; Uniprot: Q2M1Z3 / RHG31_HUMAN), S1106 and S1178, are annotated as substrates of isoform 1, whereas IV-KAPhE gives a stronger probability to isoform 2 (Figure 6a). In this case, a S6K-*α*-1 gene construct was used to induce phosphorylation under controlled conditions, while siRNAs targeting isoforms 1 and 3 were used to validate endogenous phosphorylation (Ben Djoudi Ouadda *et al*., 2018). Furthermore, site S1178 was not, in fact, tested, but rather merely identified as a putative site by S6K-*α* sequence-motif analysis (Ben Djoudi Ouadda *et al*., 2018). Finally, site T929 on protein KIBRA (*WWC1*; Uniprot: Q8IX03 / KIBRA HUMAN) is annotated to isoforms 1 and 3, however IV-KAPhE assigns low probabilities to both of these, instead favoring isoform 5 (Figure 6a). Here, the annotations are based on an *in vitro* analysis using recombinant isoforms 1 and 3 (Yang *et al*., 2014). By incorporating *in vivo* information, IV-KAPhE proposes the more likely causal isoform. I note that in many of these cases, IV-KAPhE assigns multiple kinases to the sites, albeit at varying degrees of probability, so the original assignments may be correct but incomplete.

I then carried out a similar analysis for calcium/calmodulin-dependent protein kinase type II (CaMK-II) subunits, which can form homo- or heteromultimeric holoenzymes, potentially complicating kinase assignment. As with S6K-*α*, some sites are assigned with similar probabilities to multiple subunits, while others point to possible misannotations (Supplemental Table S3; Figure 6b). For example, sites S252, S257, S282, and S285 on transcription factor C-ets-1 (*ETS1*; Uniprot: P14921 / ETS1_HUMAN) are all annotated to the most commonly studied subunit, *α*, whereas IV-KAPhE indicates that the evidence supports the least commonly studied subunit, *β* (Invergo and Beltrao, 2018), as the causal protein kinase. In the associated study, phosphorylation was tested *in vitro* using the *α* subunit, while it was tested *in vivo* through expression of a *β-γ* construct (Liu and Grundström, 2002). Interestingly, while PhosphoSitePlus only features the *in vitro* assignment to the *α* subunit, the ProtMapper corpus includes an assignment to the *β* subunit. As another example, site S2808 on ryanodine receptor 2 (RYR2; Uniprot: Q92736 / RYR2_HUMAN) is annotated to subunit *α*, whereas IV-KAPhE most strongly assigns it to the *δ* subunit (Figure 6b). In this case, the subunit or subunits used in the original, *in vitro* experiment are not specified (Rodriguez *et al*., 2003). Finally, in a similar case, sites S165 and T154 on RING finger and CHY zinc finger domain-containing protein 1 (*RCHY1*; Uniprot: Q96PM5 / ZN363 HUMAN) are annotated to subunit *α*, whereas IV-KAPhE assigns a very low probability to this subunit, strongly preferring subunit *δ*. This annotation was derived from the *in vitro* use of a CaMK-II inhibitor (Autocamtide-2 Related Inhibitory Peptide II, for which isoform specificity has not been described) and a recombinant rat kinase, the isoform of which was not specified (Duan *et al*., 2007).

### IV-KAPhE better enables high-coverage kinase activity analysis

A primary goal in performing a phosphoproteomic experiment is to deduce the signaling events that generated the data. Given quantitative phosphoproteomic measurements and a list of known substrates, the relative activity of a kinase can be inferred by a variety of methods, such as a Z-test, gene set enrichment analysis, or multiple linear regression models (Casado *et al*., 2013; Hernandez-Armenta *et al*., 2017). Typically, literature-derived kinase-substrate relationships are used, because false-positive substrate assignments from *in silico* methods introduce large variance into the pool of measurements of the substrates (Hernandez-Armenta *et al*., 2017). However, doing so inherently limits the numbers of kinases for which relative activity can be inferred.

To test whether IV-KAPhE permits more accurate inference of kinase activity over past kinase-substrate prediction or assignment methods, I performed a kinase-activity analysis on a quantitative phosphoproteomics data set in which 20 different protein kinase inhibitors were applied to MCF7 cells (Wilkes *et al*., 2015). For each condition, we expect the protein kinases targeted by the chemical inhibitor (Supplemental Table S4) to exhibit decreased activity. Furthermore, any protein kinases that are enzymatically activated by a target kinase should also exhibit decreased activity, while kinases whose activity is negatively regulated by a target kinase should show increased activity. These secondary expectations are tempered by the possibility of compensatory regulatory activity by other protein kinases. To account for these secondary inhibition effects, I identified all kinases that are likely to be regulated by each target kinase by integrating a literature-derived, signed kinase regulatory network from the Omnipath service (Türei *et al*., 2016) with a computationally predicted, signed kinase-kinase regulatory network (Invergo *et al*., 2020).

For each putatively affected kinase under each condition, I calculated Z-score-based kinase activity scores (Hernandez-Armenta *et al*., 2017), based on kinase-phosphosite assignments from each of the *in silico* methods assessed above (Figure 7a). Similarly, I calculated kinase activities using *in vivo* literature-derived annotations from PhosphoSitePlus, which provides the gold-standard in kinase activity inference (Hernandez-Armenta *et al*., 2017) (Figure 7a). Surprisingly, PhosphoSitePlus annotations enabled very few kinases to be inferred as having altered activity, however notably these were generally targets of inhibition. LinkPhinder similarly enabled few inferences of altered activity. NetworKIN 3.0, GPS 5.0, and IV-KAPhE all produced substantially more inferences, however these also included inferences of unexpected activities (e.g. up-regulation of a target kinase). Such errors could be attributable to, for example, false-positive substrate assignments, false-positive regulatory relationships, technical noise in phosphosite quantification, or compensatory phosphorylation of true-positive targets by other kinases.

**Figure 7:**
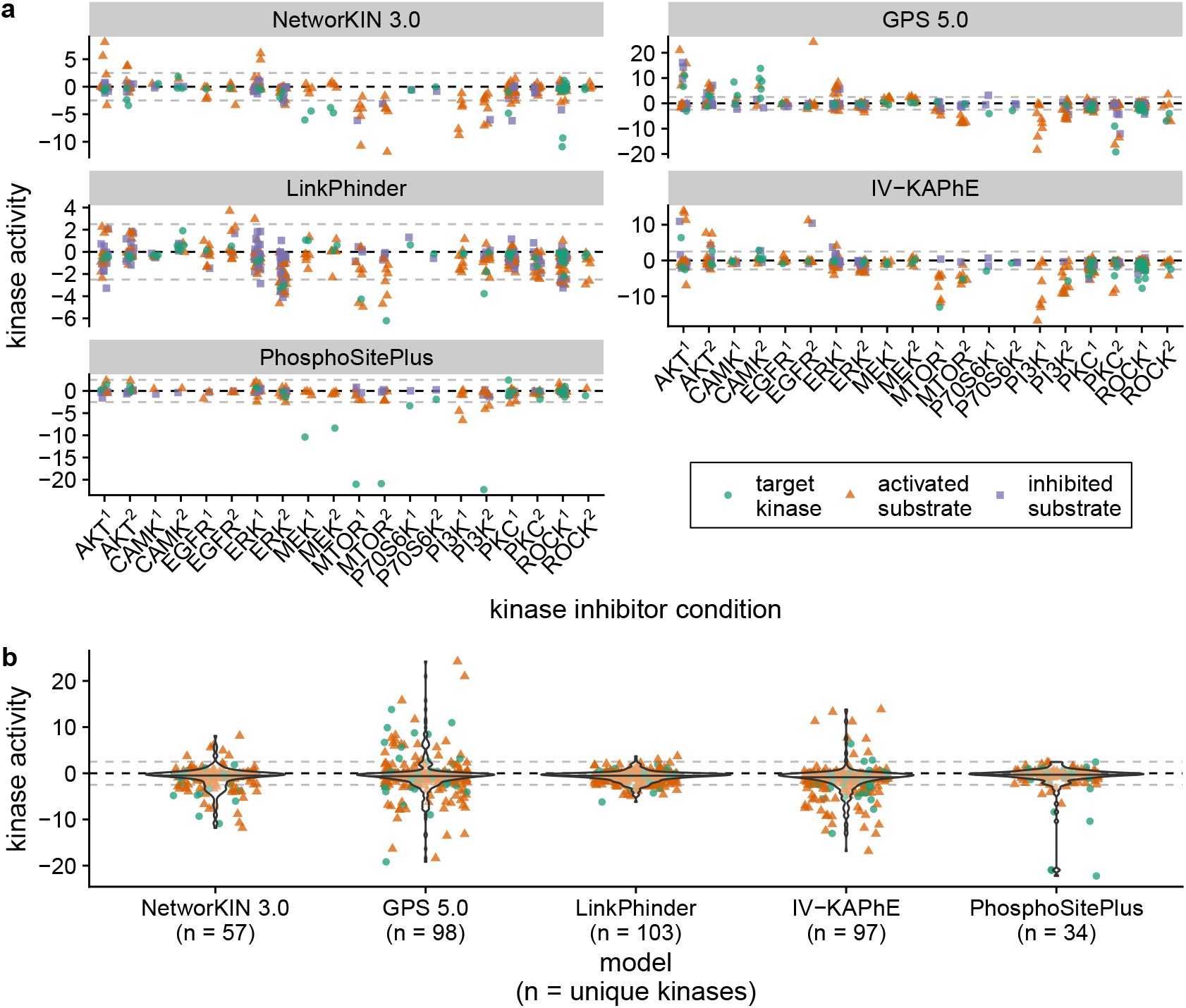
Kinase activity inference is affected by the accuracy and coverage of kinase-substrate assignments. a) Target kinases and their downstream substrate kinases in a multi-inhibitor quantitative phosphoproteomics experiment (Wilkes *et al*., 2015) are expected to have altered enzymatic activities. Assignments derived from NetworKIN 3.0, GPS 5.0, and IV-KAPhE make stronger inferences than those from LinkPhinder or *in vivo* literature-derived annotations from PhosphoSitePlus, however these methods also erroneously predict increased activity in some target kinases and downstream substrate kinases that they enzymatically activate. Each column represents a different kinase inhibitor condition (see Supplemental Table S4), in which green dots are direct targets of the inhibitor, orange triangles are kinases that are enzymatically activated by a target kinase, and violet squares are kinases that are enzymatically inhibited by a target kinase. Gray, dashed lines indicate activity levels of −2.5 and 2.5, corresponding to Z-test *p*-values of 10^−2.5^. b) IV-KAPhE provides more consistent inference of negative activity in target kinases and substrate kinases that they enzymatically activate than other computational methods as well as *in vivo* literature-derived annotations from PhosphoSitePlus. Point colors and shapes, as well as the gray dashed lines, are as described for panel (a).

Focusing on kinases that are expected to be down-regulated, ideal activity inferences would be strictly negative. By pooling these kinases from all of the conditions (noting that some kinases may appear multiple times) and observing the distribution of inferred activities, we can compare the general performance of each of the methods (Figure 7b). In pairwise comparisons, IV-KAPhE infers significantly lower activities than all of the other methods (one-sided Wilcoxon signed-rank test on matched kinase pairs, with Benjamini-Hochberg *p*-value correction for false discovery rate; IV-KAPhE vs. Networkin 3.0: *n* = 175, *V* = 4768, *p* = 6.3 × 10^−6^; vs. GPS 5.0: *n* = 262, *V* = 12272, *p* = 2.7 × 10^−5^; vs. LinkPhinder: *n* = 273, *V* = 13474, *p* = 3.1 × 10^−5^; vs. PhosphoSitePlus: *n* = 122, *V* = 1717, *p* = 1.0 × 10^−7^). The mean difference in inferred activities for matched pairs between IV-KAPhE and the other assignment sources ranged from −0.614 (LinkPhinder) to −0.943 (PhosphoSitePlus).

The kinase activity score used here is based on the *log*_10_-transformed *p*-value of the Z-test. These results indicate that the scores derived from IV-KAPhE assignments correspond to *p*-values that are, on average, up to an order of magnitude smaller than those produced from the other assignment sources. Taking appropriate precautions concerning the possibilities of false-positive assignments, then, the use of IV-KAPhE kinase-substrate assignments can provide more confident activity inferences, on average, than even *in vivo* literature-derived annotations and it does so with larger kinome coverage than literature-derived sources.

## Discussion

With the widespread availability of phosphoproteomics, methods are needed for confidently assigning protein kinases to observed phosphosites, accounting for the possibility of multiple causal kinases. Although many kinase-substrate prediction or assignment methods have been produced in the past, to the best of my knowledge no method has been specifically developed for multi-label assignment of kinases to phosphosites to meet this need. Indeed, evaluating past, performant methods in a multi-label setting, for which they were not designed, herein revealed a tendency for low average performance across their full set of covered kinases. By being built around hypothesis-free phosphoproteomic data and avoiding, where possible, functional annotations biased towards commonly studied kinases (Invergo and Beltrao, 2018), IV-KAPhE exhibits stronger average performance across the kinome, making it more suitable than past methods for the modern task of multi-label assignment of kinases to phosphoproteomic data.

Kinase-substrate annotations are not available for most species to the same level as humans or model species. Thus, kinase-substrate assignment methods that depend on such annotations cannot be applied to those species. The IV-KAPhE method presented here requires a high-throughput, phosphoproteomic kinase-substrate assay; high-throughput physical interaction data; Gene Ontology annotations, which are often have good coverage by orthology; and STRING scores, which are available for many species. This makes it a suitable method for kinase-site assignment in non-model species. While the first two requirements are, indeed, non-trivial and expensive, they are less onerous and time-consuming than low-throughput assays of individual kinase-substrates at a kinomic scale. Furthermore, the generality of the training data for the *in vivo* part of IV-KAPhE means that it may be possible to use kinase-substrate annotations from humans or model species as functional-association training data for non-model species, a possibility that remains to be explored.

Even stringently assessed, the human phosphoproteome consists of over 100,000 different sites on at least 12,000 different proteins (Ochoa *et al*., 2020), of which only a fraction have literature-derived kinase assignments. These assignments are furthermore biased towards well-studied protein kinases (Invergo and Beltrao, 2018). Thus the functional roles of many human protein kinases, including closely related isoforms of more commonly studied kinases, remain unknown. Applying IV-KAPhE predictions revealed the perils of these biases. On one hand, researchers use commonly studied isoforms for *in vitro* or artificial *in vivo* analysis, whereas the endogenous causal kinase may be a different isoform. On the other hand, when isoforms are not adequately specified in literature, annotators may default to inappropriately assigning the most commonly studied isoform to a site. As a result, the network mapping kinases to each human phosphosite remains not only largely unresolved, but its topology cannot be accurately extrapolated from existing literature-derived annotations.

Due to poor sequence conservation, many phosphosites are expected to be “off-target” and without function (Landry *et al*., 2009; Levy *et al*., 2012; Ochoa *et al*., 2020). Nevertheless, the methods presented here confidently assign multiple kinases to most sites. They collectively posit, based on the best available data, that this off-target noise is not due to low-probability events on non-optimized substrate sequences, as suggested by signaling dynamics (Kanshin *et al*., 2015). They thereby propose that the kinase-substrate network is densely connected. Incorporating further biochemical constraints into future models may reduce this apparent density and reject the computational hypothesis. Otherwise, the results will equally suggest that on-target, functional phosphorylation also can generally be catalyzed by multiple kinases, raising the question of how kinase functional specialization is maintained across the human kinome.

## Methods

### *In vitro* protein kinase specificity models

For a detailed, mathematical descriptions of specificity model construction, see Supplemental Methods. They are described here in brief. All models were built from 15-residue sequence windows around the phosphoacceptor (+/-7 residues). Substrate sequences were weighted before counting (Henikoff and Henikoff, 1994). Position-specific pseudocounts were added using the method of Henikoff and Henikoff (1996) and supplemented according to missing tail residues if sites localized to the 5’ or 3’ tail of the substrate protein. Column weights were calculated as the relative entropy versus background frequencies (PFM and phosphoproteome-backed PSSMs: position-specific residue frequencies from the full set of observed sequence windows; proteome-backed PSSMs: proteomic residue frequencies). PFM and PSSM scores were min-max normalized by calculating the theoretical minimum and maximum scores produced by the PSSM (Wagih *et al*., 2015).

The multi-label Naïve Bayes model was built using four components: the PFM constructed from a kinase’s substrates; the PFM of all the substrates not phosphorylated by the kinase; and the prior probabilities of a site being phosphorylated by the kinase or by any other kinase. The priors were empirically estimated as the fraction of substrates in the experiment that were phosphorylated by the kinase or not, respectively. For determining the posterior probability that a kinase *K* phosphorylates a site with sequence window **S**, given PFM scores *s_K_* for *K* and 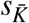 for all other kinases and prior probabilities *P*(*K*) and 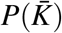, the following was calculated:

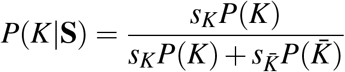

A kinase was assigned to a site if *P*(*K*|**S**) > 0.5.

The Naïve Bayes+ model further incorporated Bernoulli-distributed features with the PFM-based likelihood function. Probability parameters of substrates being direct or indirect interaction partners, of carrying domains enriched among a kinase’s substrates or interaction partners, or of substrate sites being within a predicted protein domain were estimated empirically from the training data. Domain enrichment was calculated among substrates or interaction partners via Fisher’s hypergeometric test and *p*-values were adjusted for false-discovery rate before being tested at a critical value of 0.05. To facilitate the scoring of new human phosphosites, the Naïve Bayes+ model for human kinases, as generated and applicable by the motif-kit software (see “Implementation”, below), as well as the domain enrichment and kinase interaction data have been archived on Zenodo (doi: 10.5281/zenodo.6325198).

### Multi-label cross-validation

For multi-label performance evaluation, I performed 10 iterations of 10-fold cross-validation, restricting to kinases with models trained on at least 20 substrates. Both kinase substrate-domain enrichment and sequence specificity models were recalculated from each fold’s training subset. Performance was evaluated using macro-averaged (averaged over the set of all kinases, 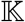) precision, recall and F1 scores:

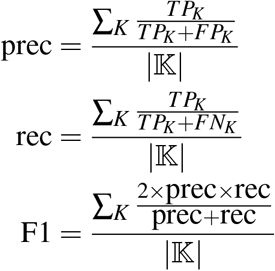

where *TP_K_*, *FP_K_*, and *FN_K_* are the true-positive, false-positive, and false-negative counts for kinase *K*, respectively. If a score was undefined due to division by 0, it was set to 0.

### Implementation

Kinase specificity model training and scoring were implemented in and performed using a bespoke, free and open-source toolkit called *motif-kit* (https://www.gitlab.com/brandoninvergo/motif-kit). Version 1.0, used here, is archived at Zenodo (doi: 10.5281/zenodo.6325136). The code was written in ANSI C99 for POSIX systems and is dependent only on the GNU Scientific Library (https://www.gnu.org/software/gsl) and the HDF5 library (https://www.hdfgroup.org/solutions/hdf5/), with unit tests further depending on the Check testing framework (https://libcheck.github.io/check/). All other methods (domain enrichment, multi-label CV, etc.) were implemented using R (version 4.1.0).

### IV-KAPhE Model

#### Model construction

The IV-KAPhE model was built via the Random Forest method, as implemented in the R package “ranger” (version 0.13.1; Wright and Ziegler, 2017). The models were built with 500 trees. The variable importance mode (parameter “importance”) was set to “impurity” (the Gini index) and the splitting rule (parameter “splitrule”) was set to “gini”. The model was trained to classify a given kinase-phosphosite pair as “true” (the kinase phosphorylates the site) or “false” (it does not) and set to provide a probability for each class. Feature selection was performed via variable-importance analysis, as implemented in ranger. The final list of features retained for model construction were: Naïve Bayes+ posterior probability, GO BP and CC semantic similarity, STRING coexpression and experimental scores, whether the substrate protein is itself a kinase, the kinase type (S/T or Y), and the site class (S/T or Y). The model was also tested using a Support Vector Machine instead of Random Forest, as implemented in the R package “e1071” (version 1.7-9) using the default settings for classification.

#### Cross-validation

*In vivo* kinase-substrate relationships identified in the PhosphoSitePlus database (Hornbeck *et al*., 2015) were used as true positives in the training set for cross-validation. The same kinases from the true positive set (including multiple occurrences) were randomly assigned to other sites from the human phosphoproteome (Ochoa *et al*., 2020) to form a negative set. To this end, S/T kinases were randomly assigned to S/T sites and Y kinases were assigned to Y sites. Additionally, any S/T or Y kinases that were annotated as phosphorylating the opposite site type in the true positive set were randomly assigned to a similar proportion of such sites in the negative set. In all cases, the proportion of substrate kinases observed in the positive set was maintained in the random negative set. Sites were filtered not to include sites found in the true positive training set or the external testing set (see below). 10-fold cross validation was performed and evaluated, restricted to kinases with Naive Bayes+ models trained on at least 20 substrates, via multi-label precision, recall, and F1 as described above. Cross-validation was performed ensuring that any kinase present in a fold had at least one positive and one negative site.

#### External Validation

IV-KAPhE was trained using the full PhosphoSitePlus and random kinase-site pair training set as described above. An external evaluation set was prepared by identifying kinase-substrate relationships inferred via the ProtMapper method (Bachman *et al*., 2019) which were not present in PhosphoSitePlus (*in vivo* or *in vitro*). These sites were accompanied by further random negative kinase-site pairs as described above. Predictions made by the model on this testing set were evaluated via multi-label precision, recall, and F1.

I furthermore evaluated the assignments for these kinase-site pairs made by phosphoproteome-backed PSSMs, Naïve Bayes+, and three other, previously published tools with similar kinomic scope or model architecture: NetworKIN 3.0 (Horn *et al*., 2014), GPS 5.0 (Wang *et al*., 2020), and LinkPhinder (Nováček *et al*., 2020). NetworKIN and GPS were run in-house with their default settings, whereas the LinkPhinder scores produced by the authors were used. For further evaluation, the published stringent LinkPhinder cutoff of 0.672 (Nováček *et al*., 2020) and the nominal NetworKIN cutoff of 1.0 (Horn *et al*., 2014) were used.

The test set was then re-evaluated with all methods, now allowing all-versus-all kinase-site assignments, restricted to sites that had at least one true kinase assigned. The rate of novel assignments was estimated via the macro-averaged false discovery rate (FDR) and compared to the macro-averaged true positive rate (identical to recall):

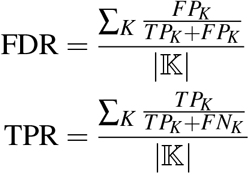

“False positive” (FP) is used here by convention although it is a misnomer in the present case, as we do not know that the assignments are false.

### Kinase Assignment Distributions

IV-KAPhE and Naïve Bayes+ were applied to perform all-versus-all assignments for kinases with specificity models built from at least 20 substrates against the full high-confidence set of human phosphosites (Ochoa *et al*., 2020). The numbers of kinases assigned to each phosphosite and the median number of kinases assigned to sites on each substrate protein were analyzed via histograms. All literature-derived assignments in PhosphoSitePlus and all assignments provided by the authors of LinkPhinder at a cutoff of 0.672 were analyzed similarly.

### Kinase Activity Analysis

Previously published quantitative phosphoproteomic measurements from a multi-inhibitor experiment (Wilkes *et al*., 2015) were filtered to remove missing data and measurements taken under serine/threonine-protein phosphatase 2A inhibition. If a phosphosite was observed on multiple peptides in the data, the peptide with the greatest dynamic range between conditions was retained. Protein kinases that are regulated downstream of the kinases targeted for inhibition were retrieved from two sources: Omnipath, a meta-database of protein-protein regulatory relationships (Türei *et al*., 2016), keeping only kinase-kinase regulatory relationships with a consensus sign (activating or inhibiting); and a set of computational predictions of signed kinase-kinase regulatory relationships (Invergo *et al*., 2020), with a stringent posterior-probability cutoff of 0.75. Multi-label kinase-substrate assignment was then performed for all target kinases and each of their regulatory-substrate kinases using IV-KAPhE, NetworKIN 3.0, and LinkPhinder at their nominal cutoffs as described above and using GPS 5.0 at its “medium” stringency setting. Furthermore, *in vivo* kinase-substrate annotations were retrieved from PhosphoSitePlus for the sites.

For each kinase-substrate assignment source, kinase activities were inferred as follows. For a given kinase and inhibition condition, the *log*_2_ fold-changes of any of the kinase’s assigned substrates were tested for significant difference from the mean *log*_2_ fold-change for that condition via a two-sided Z-test. The final activity was inferred as 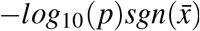, where *sgn* is the sign function, 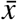 is the mean fold-change of the kinase’s assigned substrates, and *p* is the *p*-value of the Z-test. For example, if the kinase’s assigned substrates have a significantly lower distribution than the full sample, the inferred activity will take a large negative value.

## Acknowledgments

This work was supported by a Wellcome Trust Institutional Strategic Support Award [204904/Z/16/Z]. For the purpose of open access, the author has applied a CC BY public copyright licence to any Author Accepted Manuscript version arising from this submission. The author would like to thank Pedro Beltrao for helpful comments on a draft of this manuscript.

