## Supplemental Materials for "Accurate, high-coverage assignment of *in vivo* protein kinases to phosphosites from *in vitro* phosphoproteomic specificity data"

Brandon M. Invergo[1,*]

[1]Translational Research Exchange @ Exeter, University of Exeter, Stocker Road, Exeter EX4 4QJ

### Supplemental Methods

#### Data

*In vitro* kinase substrates were retrieved from Sugiyama *et al.* (2019), excluding mutant kinases and lipid kinases. Autophosphorylation events were omitted from the analysis. In order to stringently control for false positives, kinase substrates were filtered to include only those that passed both the authors' "SIDIC" and "PTMscore" ($P > 0.75$) criteria (Sugiyama *et al.*, 2019). A high-confidence set of human phosphosites was retrieved from Ochoa *et al.* (2020). Literature-derived, human phosphosites and kinase-substrate relationships were retrieved from the PhosphoSitePlus database (version 020922; Hornbeck *et al.*, 2015) and from Bachman *et al.* (2019), and kinase-substrate relationships filtered to remove autophosphorylation.

Kinase physical and indirect interactions were derived from a merged network of interactions from the BioGRID database (version 4.3.194; Oughtred *et al.*, 2021), the BioPlex 293T network (December 2019; Huttlin *et al.*, 2021), and a mass-sepctrometry-based assay of kinase interactions (Buljan *et al.*, 2020). BioGRID interactions were filtered to exclude biochemical reactions ("Experimental System" = "Biochemical Activity") and genetic interactions. All networks were filtered to exclude self-binding. Protein domains were retrieved from Pfam (version 33.1; Finn *et al.*, 2016). Protein functional association scores were retrieved from the STRING database (version 11.0; Szklarczyk *et al.*, 2019). Kinase identities (for IV-KAPhE "substrate is a kinase" feature) and types (S/T or Y) were derived from the *pkinfam.txt* file provided by UniProt and Swiss-Prot (release 2020_02; `https://www.uniprot.org/docs/pkinfam`). For this purpose, dual-specificity kinases were counted as S/T kinases.

##### Gene Ontology semantic similarity

Semantic similarity of Gene Ontology (GO) terms between a kinase and a putative substrate protein was calculated using the "GOSemSim" package for R (version 2.18.0; Yu *et al.*, 2010; Yu, 2020). Semantic similarity was calculated using the "Wang" measure and combined using the "BMA" method. Missing values (due to lack of annotation) were imputed from the full *in vivo* feature table via multivariate imputation by chained equations using the "mice" package for R (40 maximum iterations; version 3.13.0; van Buuren and Groothuis-Oudshoorn, 2011).

##### STRING functional association scores

For all STRING component scores considered ("coexpression", "experimental", "cooccurence", "fusion", and "neighborhood"), values were transformed to be between 0 and 1 by dividing by 1000. Missing values were set to the prior value of 0.041 used by the STRING developers. STRING uses Ensemble identifiers for proteins, whereas the rest of the analysis used Uniprot identifiers. In some cases, many Ensembl identifiers can map to the same Uniprot identifier. In these cases, the means of the scores assigned to all relevant identifiers were used.

#### Kinase specificity models

##### Position Frequency Matrix

Sequence windows were first weighted using a position-based sequence-weighting method (Henikoff and Henikoff, 1994). Given a set of $n \geq 20$ substrate sequence windows, $\mathbb{S} = \mathbf{S}_1, \mathbf{S}_2, \ldots, \mathbf{S}_i, \ldots, \mathbf{S}_{n-1}, \mathbf{S}_n$, where $\mathbf{S}_i = S_{i1}, S_{i2}, \ldots, S_{i14}, S_{i15}$ and $S_{ij}$ represents the amino acid at position $j$ of sequence $i$, a weight was assigned to each amino acid $a$ at position $j$:

$$w(a, j) = \frac{1}{c_j \sum_{i=1}^{n} (S_{ij} = a)} \tag{1}$$

where $c_j$ is the number of unique amino acids found at position $j$ among the substrates in $\mathbb{S}$. Phosphosites near the 5' or 3' tail of the protein may not have 7 residues adjacent to them to form a complete sequence window. At each missing tail position, a $21^{\text{st}}$ amino acid was counted. A weight was calculated for each sequence as the sum of its position-specific residue weights:

$$W(\mathbf{S}_i) = \sum_{j=1}^{15} w(S_{ij}, j) \tag{2}$$

Finally, each sequence's weight was normalized:

$$\widehat{W}(\mathbf{S}_i) = \frac{W(\mathbf{S}_i)}{\sum_{k=1}^{n} W(\mathbf{S}_k)} \tag{3}$$

A $20 \times 15$ counts matrix, $\mathbf{r}$ was constructed such that entry $r_{aj}$ contains the weighted count of amino acid $a$, ignoring missing tails, at position $j$ across the sequences in $\mathbb{S}$:

$$r_{a,j} = n \sum_{i=1}^{n} V(S_{ij}, a) \tag{4}$$

$$V(S_{ij}, a) = \begin{cases} \widehat{W}(\mathbf{S}_i) & \text{if } S_{ij} = a \\ 0 & \text{otherwise} \end{cases} \tag{5}$$

To account for the fact that $\mathbb{S}$ necessarily represents a limited sample of possible substrate sequences, a total number of pseudocounts, $B_j$, were distributed at position $j$ via $20 \times 15$ pseudocount matrix $\mathbf{b}$ as described below. The final PFM, $\mathbf{p}$ was then calculated from the counts and pseudocounts such that the likelihood of observing amino acid $a$ at position $j$ is:

$$PFM_{a,j} = p_{a,j} = \frac{b_{a,j} + r_{a,j}}{B_j + \sum_a r_{a,j}} \tag{6}$$

Following Henikoff and Henikoff (1996), a position-specific pseudocount strategy was adopted. For column $j$ the total number of pseudocounts to be distributed among the amino acids is:

$$B_j = T_j + m \times c_j \tag{7}$$

where $T_j$ is the number of missing tails observed in position $j$ among the substrates in $\mathbb{S}$, $c_j$ is defined as in Equation 1, and $m$ is a tune-able parameter, which was fixed at 1 (Invergo *et al.*, 2020). Note that positions with lower diversity of observed residues will receive fewer pseudocounts.

Pseudocounts were then distributed according to a $20 \times 15$ expected frequency matrix $\mathbf{f}$. In the case of PFMs evaluated directly for predictive performance or used in the construction of phosphoproteome-backed PSSMs, $\mathbf{f}$ was constructed as a PFM (as above) on the full set of *in vitro* substrates from all kinases, $\mathbb{S}_{BG}$, with no pseudocounts ($B_j = 0$). In the case of PFMs used in the construction of proteome-backed PSSMs, $\mathbf{f}$ was constructed such that element $f_{a,j} = F_a$, where $F_a$ is the proteomic frequency of residue $a$. Pseudocount matrix $\mathbf{b}$ was then calculated such that residue $a$ at position $j$ receives $b_{a,j}$ pseudocounts:

$$b_{a,j} = B_j \times f_{a,j} \tag{8}$$

Finally, for each position $j$, a column weight $\hat{w}_j$ was calculated as the relative entropy (Kullback-Leibler divergence) of the residue frequencies at that position, $\mathbf{p}_j$, versus the background expectation, $\mathbf{f}_j$, rescaled to sum to 15 (the number of positions):

$$w_j = \sum_a p_{a,j} \log_2 \left( \frac{p_{a,j}}{f_{a,j}} \right) \tag{9}$$

$$\hat{w}_j = w_j \times \frac{15}{\sum_j w_j} \tag{10}$$

To assign an unseen substrate, $\mathbf{S}$, to a kinase with PFM $\mathbf{p}$ and column weights $\hat{\mathbf{w}}$, a score is calculated as:

$$s_{PFM}(\mathbf{S}, \mathbf{p}, \hat{\mathbf{w}}) = \prod_j \hat{w}_j p_{S_j,j} \tag{11}$$

**Position-Specific Scoring Matrix**

$20 \times 15$ PSSMs were calculated as log-likelihood ratios against the background expectation $\mathbf{f}$:

$$PSSM_{a,j} = \hat{p}_{a,j} = \log_2 \left( \frac{p_{a,j}}{f_{a,j}} \right) \tag{12}$$

As described above, "proteome-backed" PSSMs were calculated against a background expectation of proteomic residue frequencies, while "phosphoproteome-backed" PSSMs were calculated against a position-specific background expectation derived from all phosphosite sequences observed in the experiment.

For assigning an unseen substrate, $\mathbf{S}$, to a kinase with PSSM $\hat{\mathbf{p}}$ and column weights $\hat{\mathbf{w}}$, a score is calculated as:

$$s_{PSSM}(\mathbf{S}, \hat{\mathbf{p}}, \hat{\mathbf{w}}) = \sum_j \hat{w}_j \hat{p}_{S_j,j} \tag{13}$$

For multi-label assignment, a min-max normalized PSSM score, $\hat{s}_{PSSM}$, was calculated to take values between 0 and 1 (Wagih *et al.*, 2015):

$$a_{min} = \arg\min_a \hat{p}(a, j) \tag{14}$$

$$a_{max} = \arg\max_a \hat{p}(a, j) \tag{15}$$

$$s_{min} = \sum_j \hat{w}_j \hat{p}(a_{min}, j) \tag{16}$$

$$s_{max} = \sum_j \hat{w}_j \hat{p}(a_{max}, j) \tag{17}$$

$$\hat{s}_{PSSM} = \frac{s_{PSSM} - s_{min}}{s_{max} - s_{min}} \tag{18}$$

**Multi-label Naïve Bayes**

We wish to calculate the posterior probability that a phosphosite is phosphorylated by kinase $K$, given its surrounding sequence window $\mathbf{S}$. Using Bayes' Theorem:

$$P(K|\mathbf{S}) = \frac{\mathscr{L}(\mathbf{S}|K)P(K)}{P(\mathbf{S})} \tag{19}$$

$$= \frac{\mathscr{L}(S_1, S_2, \ldots, S_{14}, S_{15}|K)P(K)}{P(\mathbf{S})} \tag{20}$$

By applying the chain rule and then the Naïve Bayes assumption of conditional independence between the positions in **S**, this can be simplified as follows:

$$P(K|\mathbf{S}) = \frac{\mathscr{L}(S_1|S_2, \ldots, S_{15}, K)\mathscr{L}(S_2|S_3, \ldots, S_{15}, K)\ldots\mathscr{L}(S_{14}|S_{15}, K)\mathscr{L}(S_{15}|K)P(K)}{P(\mathbf{S})} \tag{21}$$

$$= \frac{\mathscr{L}(S_1|K)\mathscr{L}(S_2|K)\ldots\mathscr{L}(S_{14}|K)\mathscr{L}(S_{15}|K)P(K)}{P(\mathbf{S})} \tag{22}$$

$$= \frac{P(K)\prod_j \mathscr{L}(S_j|K)}{P(\mathbf{S})} \tag{23}$$

We model the likelihood function, $\mathscr{L}(S_j|K)$ as a categorical distribution. It is then easy to see from Equation 11 that, with the Naïve Bayes assumption of conditional independence:

$$\mathscr{L}(\mathbf{S}|K) = \prod_j \mathscr{L}(S_j|K) = s_{PFM}(\mathbf{S}, \mathbf{p}_K, \hat{\mathbf{w}} = \{1, 1, \ldots, 1, 1\}) \tag{24}$$

By altering the values in $\hat{\mathbf{w}}$, we arrive at a weighted Naïve Bayes formulation:

$$P(K|\mathbf{S}) = \frac{P(K)\prod_j \hat{w}_j \mathscr{L}(S_j|K)}{P(\mathbf{S})} \tag{25}$$

$$= \frac{P(K)s_{PFM}(\mathbf{S}, \mathbf{p}_K, \hat{\mathbf{w}}_K)}{P(\mathbf{S})} \tag{26}$$

So, the conditional likelihood function $\mathscr{L}(\mathbf{S}|K)$ is calculated as described for PFMs.

The prior probability, $P(K)$ was empirically estimated as the proportion of all substrates observed in the experiment ($\mathbb{S}_{BG}$) that were observed as phosphorylated by kinase $K$ ($\mathbb{S}_K$):

$$P(K) = \frac{|\mathbb{S}_K|}{|\mathbb{S}_{BG}|} \tag{27}$$

Typically the marginal probability, $P(\mathbf{S})$, is ignored in Naïve Bayes assignment as it is just a normalizing constant. However, in multi-label Naïve Bayes, normalization is necessary to compare probabilities between classes (kinases) (Zhang *et al.*, 2009). By the law of total probability:

$$P(\mathbf{S}) = \mathscr{L}(\mathbf{S}|K)P(K) + \mathscr{L}(\mathbf{S}|\bar{K})P(\bar{K}) \tag{28}$$

The second term represents the likelihood of sequence **S** being phosphorylated by any other kinase in the experiment ($\bar{K}$). This is computed exactly as for kinase $K$, using sequences not phosphorylated by $K$, $\mathbb{S}_{\bar{K}}$, to build the PFM and calculate the prior probability:

$$\mathbb{S}_{\bar{K}} = \mathbb{S}_{BG} - \mathbb{S}_K \tag{29}$$

To perform multi-label assignment of phosphosite sequence **S**, for each kinase $K_i$ in the set of kinases used in the experiment, $\mathbb{K}$, posterior probabilities were calculated as

$$P(K_i|\mathbf{S}) = \frac{P(K_i)s_{PFM}(\mathbf{S}, \mathbf{p}_{K_i}, \hat{\mathbf{w}}_{K_i})}{P(K_i)s_{PFM}(\mathbf{S}, \mathbf{p}_{K_i}, \hat{\mathbf{w}}_{K_i}) + P(\bar{K}_i)s_{PFM}(\mathbf{S}, \mathbf{p}_{\bar{K}_i}, \hat{\mathbf{w}}_{\bar{K}_i})} \tag{30}$$

$$P(\bar{K}_i|\mathbf{S}) = \frac{P(\bar{K}_i)s_{PFM}(\mathbf{S}, \mathbf{p}_{\bar{K}_i}, \hat{\mathbf{w}}_{\bar{k}_i})}{P(K_i)s_{PFM}(\mathbf{S}, \mathbf{p}_{K_i}, \hat{\mathbf{w}}_{K_i}) + P(\bar{K}_i)s_{PFM}(\mathbf{S}, \mathbf{p}_{\bar{K}_i}, \hat{\mathbf{w}}_{\bar{K}_i})} \tag{31}$$

Kinase $K_i$ was assigned to the phosphosite with sequence **S** if $P(K_i|\mathbf{S}) > P(\bar{K}_i|\mathbf{S})$ (Zhang *et al.*, 2009). This is equivalent to the condition that $P(K_i|\mathbf{S}) > 0.5$, thus $P(\bar{K}_i|\mathbf{S})$ need not be calculated.

**Naïve Bayes+**

The Naïve Bayes+ model is an extension of the multi-label Naïve Bayes model described above. In addition to the sequence evidence **S**, additional Bernoulli-distributed features were added – $I_{K,T}$ or $\bar{I}_{K,T}$: presence or absence of a direct physical interaction between kinase $K$ and substrate protein $T$; $J_{K,T}$ or $\bar{J}_{K,T}$: presence or absence of an indirect physical interaction between them; $D$ or $\bar{D}$: the phosphosite is or is not in a predicted domain on $T$; and $E$ or $\bar{E}$: substrate $T$ does or does not carry a predicted domain that is enriched among $K$'s substrates or physical interaction partners. An indirect interaction was defined as the substrate protein $T$ being "two hops" away from $K$ on the protein interaction network, i.e. they share a neighbor.

Thus, for example, the Naïve Bayes formulation of the posterior probability that kinase $K$ phosphorylates a site with sequence **S**, that is not in a domain $\bar{D}$, with no direct physical interaction $\bar{I}_{K,T}$, an indirect interaction $J_{K,T}$, and no enriched domains $\bar{E}$ would be:

$$P(K|\mathbf{S}, \bar{D}, \bar{I}_{K,T}, J_{K,T}, \bar{E}) = \frac{\mathscr{L}(\mathbf{S}|K)\mathscr{L}(\bar{D}|K)\mathscr{L}(\bar{I}_{K,T}|K)\mathscr{L}(J_{K,T}|K)\mathscr{L}(\bar{E}|K)P(K)}{P(S)} \tag{32}$$

$$\begin{aligned} P(S) = {} & \mathscr{L}(\mathbf{S}|K)\mathscr{L}(\bar{D}|K)\mathscr{L}(\bar{I}_{K,T}|K)\mathscr{L}(J_{K,T}|K)\mathscr{L}(\bar{E}|K)P(K) + \\ & \mathscr{L}(\mathbf{S}|\bar{K})\mathscr{L}(\bar{D}|\bar{K})\mathscr{L}(\bar{I}_{K,T}|\bar{K})\mathscr{L}(J_{K,T}|\bar{K})\mathscr{L}(\bar{E}|\bar{K})P(\bar{K}) \end{aligned} \tag{33}$$

For all four additional features, the Bernoulli parameter $p$ was estimated by calculating the fraction of $K$'s substrates that have the feature (or, for $\bar{K}$, the fraction of all sites that are not substrates of $K$). Domains enriched within the substrate proteins or interaction partners of kinase $K$ were identified via Fisher's Hypergeometric Test. For each unique domain predicted to be on at least one substrate protein, presence or absence of that domain among proteins were counted contingent upon the proteins being either substrates or physical interaction partners of $K$ or not. $p$-values were adjusted for false discovery rate and tested at a critical value of 0.05.

#### Approximate logistic relationship between phosphoproteome-backed PSSM score and Naïve Bayes posterior probability

As described above, the definition of the Naïve Bayes posterior probability of assigning kinase $K$ to a phosphosite with sequence window $\mathbf{S}$ is:

$$P(K|\mathbf{S}) = \frac{\mathscr{L}(\mathbf{S}|K)P(K)}{P(\mathbf{S})} \tag{34}$$

$$\mathscr{L}(\mathbf{S}|K) = \mathscr{L}(S_1|K)\mathscr{L}(S_2|K)...\mathscr{L}(S_{14}|K)\mathscr{L}(S_{15}|K) \tag{35}$$

In terms of the likelihood, the phosphoproteome-backed PSSM score function $s_{PSSM}(\mathbf{S}, \hat{\mathbf{p}}_K, \hat{\mathbf{w}}_K)$ can be re-written as:

$$s_{PSSM}(\mathbf{S}, \hat{\mathbf{p}}_K, \hat{\mathbf{w}}_K) = \log_2\left(\frac{\mathscr{L}(\mathbf{S}|K)}{\mathscr{L}(\mathbf{S})}\right) \tag{36}$$

Likelihood $\mathscr{L}(\mathbf{S})$ is the overall likelihood of observing a phosphosite with sequence window $\mathbf{S}$, as calculated using the full phosphoproteome background set, $\mathbb{S}_{BG}$. Likelihood $\mathscr{L}(\mathbf{S}|K)$ is the likelihood of observing a phosphosite with sequence window $\mathbf{S}$ among the substrates of kinase $K$, as calculated using a set of known substrates of kinase $K$, $\mathbb{S}_K$. Furthermore, we assume:

$$\mathbb{S}_K \subset \mathbb{S}_{BG} \tag{37}$$

$$|\mathbb{S}_K| \ll |\mathbb{S}_{BG}| \tag{38}$$

We apply a logit transformation to the Naïve Bayes posterior probability $P(K|\mathbf{s})$, using 2 as our logarithm base:

$$\begin{aligned}
\text{logit}(P(K|\mathbf{S})) &= \log_2\left(\frac{P(K|\mathbf{S})}{1-P(K|\mathbf{S})}\right) \\
&= \log_2\left(\frac{P(K|\mathbf{S})}{P(\overline{K}|\mathbf{S})}\right) \\
&= \log_2\left(\frac{\mathscr{L}(\mathbf{S}|K)P(K)}{\mathscr{L}(\mathbf{S}|\overline{K})P(\overline{K})}\right) \\
&= \log_2\left(\frac{\mathscr{L}(\mathbf{S}|K)}{\mathscr{L}(\mathbf{S}|\overline{K}))}\right) + \log_2\left(\frac{P(K)}{P(\overline{K})}\right)
\end{aligned} \tag{39}$$

Given that $|\mathbb{S}_K| \ll |\mathbb{S}_{BG}|$ (Equation 38), $\mathscr{L}(\mathbf{S}|\overline{K}) \approx \mathscr{L}(\mathbf{S})$. Put simply, the likelihood model constructed from a set of over 14000 phosphosites will not be strongly impacted if one were first to omit a subset of a few hundred sites on average. As a result:

$$\begin{aligned}
\mathrm{logit}(P(K|\mathbf{S})) &\approx \log_2\left(\frac{\mathscr{L}(\mathbf{S}|K)}{\mathscr{L}(\mathbf{S}))}\right) + \log_2\left(\frac{P(K)}{P(\bar{K})}\right) \\
\mathrm{logit}(P(K|\mathbf{S})) &\approx s_{PSSM}(\mathbf{S}, \hat{\mathbf{p}}_K, \hat{\mathbf{w}}_K) + \log_2\left(\frac{P(K)}{P(\bar{K})}\right) \\
s_{PSSM}(\mathbf{S}, \hat{\mathbf{p}}_K, \hat{\mathbf{w}}_K) &\approx \mathrm{logit}(P(K|\mathbf{S})) - \log_2\left(\frac{P(K)}{P(\bar{K})}\right)
\end{aligned} \tag{40}$$

From Equation 38, it also follows that $|\mathbb{S}_K| < |\mathbb{S}_{\bar{K}}|$, so $P(K) < P(\bar{K})$ and $\log_2\left(\frac{P(K)}{P(\bar{K})}\right) < 0$. This imposes a dependence of the relationship on $|\mathbb{S}_K|$: a larger $|\mathbb{S}_K|$ causes a lower inflection point in the inverse logistic fit, i.e. the PSSM score $s_{PSSM}(\mathbf{S}, \hat{\mathbf{p}}_K, \hat{\mathbf{w}}_K)$ at which $P(K|\mathbf{S}) = 0.5$ and $\mathrm{logit}(P(K|\mathbf{S})) = 0$.

### Supplemental Figures

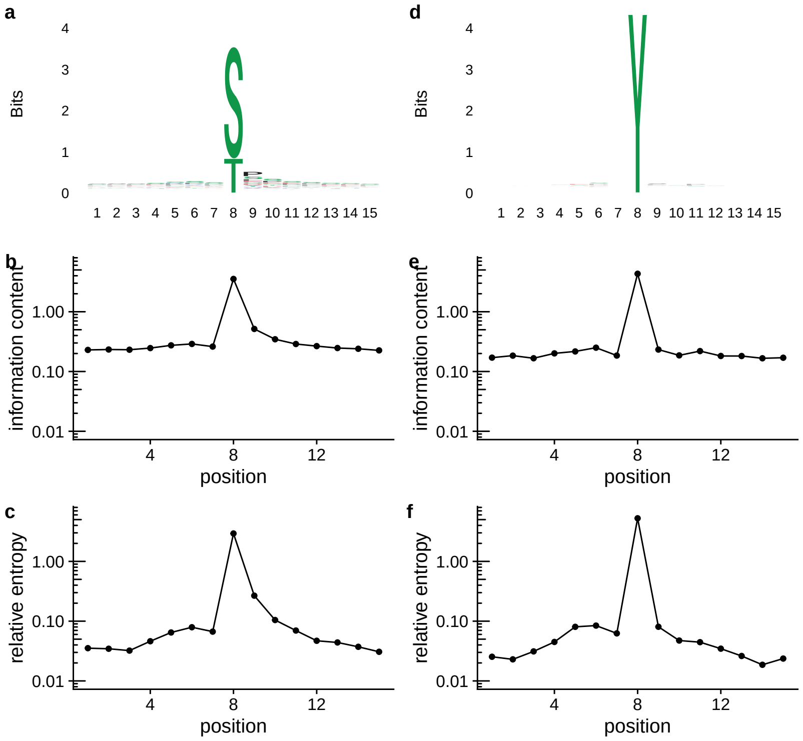


Figure S1: a) Sequence logo of all S/T sites in a high-confidence human phosphoproteome (Ochoa *et al.*, 2020). b) Information content at each position among S/T sites. c) Relative entropy versus proteomic residue frequencies at each position among S/T sites. Relative entropy provides greater separation between positions. d-f) As in (a-c) for Y sites.

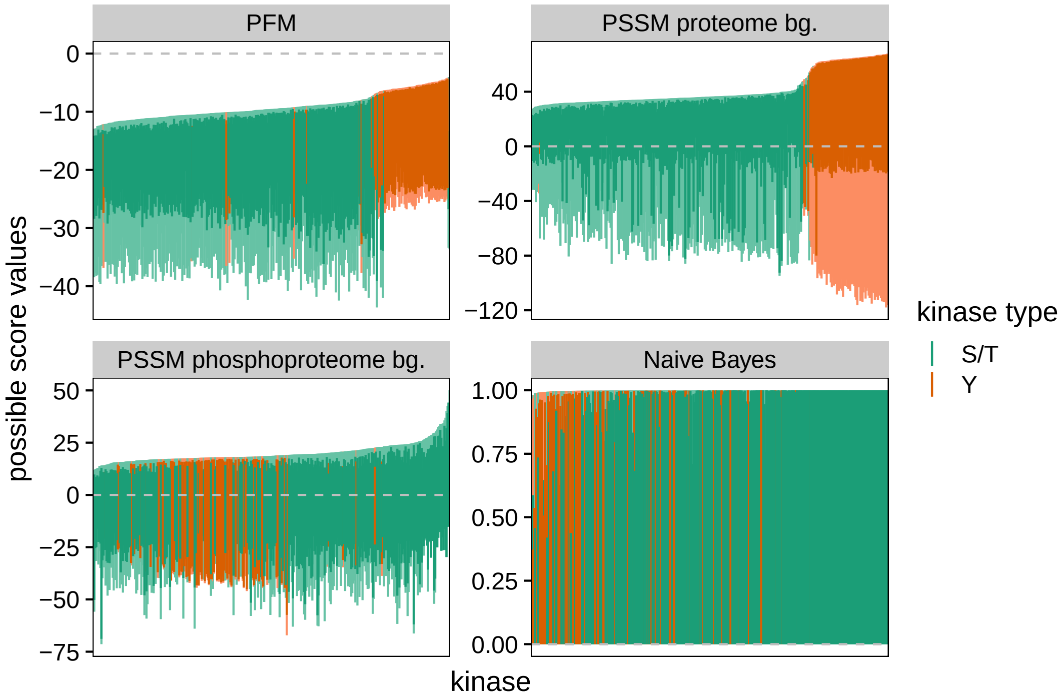


Figure S2: Theoretical (light colors) and empirical (dark colors) score ranges for the four sequence-only kinase specificity models. Empirical ranges were identified using the high-confidence human phosphoproteome (Ochoa *et al.*, 2020). PFM scores are shown $\log_{10}$-transformed. Each vertical line represents one kinase, colored by S/T (green) or Y kinases (orange). Note that PFM and PSSM-based models have very different ranges between kinases. The maximum theoretical also varies, making universal cutoff definitions difficult. Also, PFMs and PSSMs do not utilize their full range for all kinases when scoring physiologically occurring phosphosites. The Naïve Bayes model exhibits none of these problems.

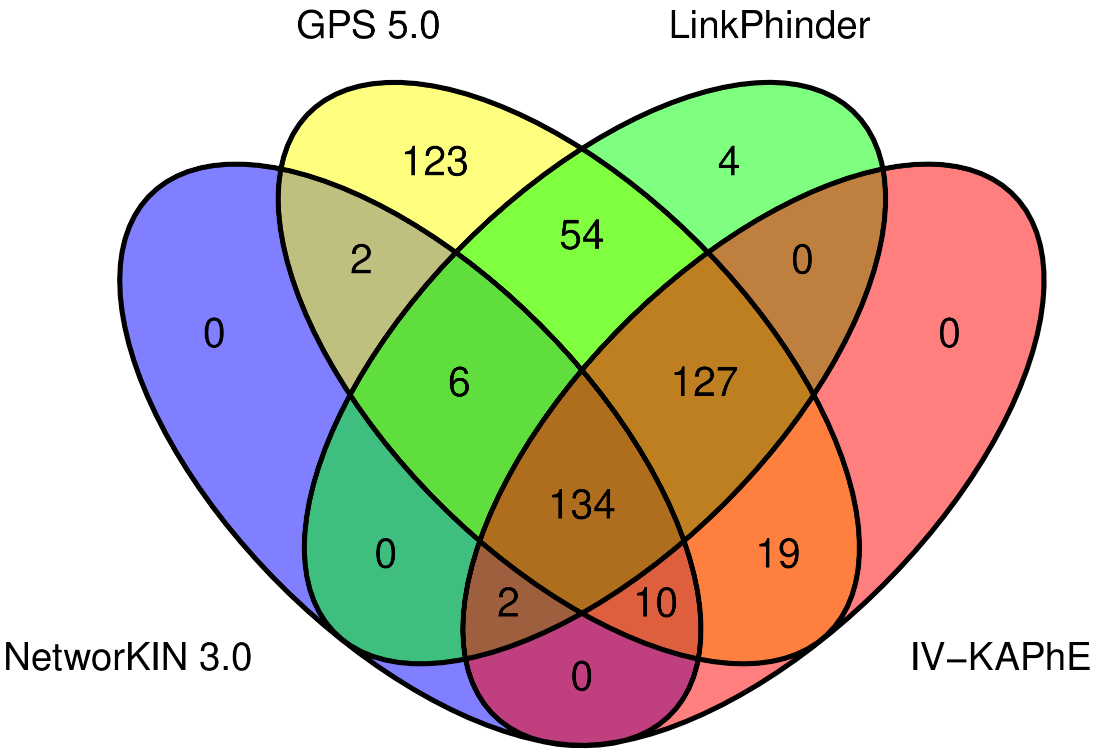


Figure S3: A Venn diagram showing total coverage of the human kinome supported by four kinase-substrate prediction or assignment methods. Values reflect numbers of kinases. GPS 5.0 has the largest coverage, while NetworKIN 3.0's coverage is significantly lower than the others. IV-KAPhE does not cover any kinases not covered by the other methods.

### Supplemental Tables

Supplemental Table S1 (TSV File): All-versus-all, kinase-substrate assignments for a high-confidence human phosphoproteome (Ochoa *et al.*, 2020) calculated using Näive Bayes+ and IV-KAPhE.

—

Supplemental Table S2 (TSV File): IV-KAPhE assignments of S6K-$\alpha$ isoforms for any site annotated in PhosphoSitePlus as being phosphorylated by at least one of these isoforms *in vivo* or *in vitro*.

—

Supplemental Table S3 (TSV File): IV-KAPhE assignments of CaMK-II subunits for any site annotated in PhosphoSitePlus as being phosphorylated by at least one of these subunits *in vivo* or *in vitro*.

—

Supplemental Table S4: Protein kinases targeted by the chemical inhibitors used in the quantitative phosphoproteomic experiment of Wilkes *et al.* (2015).

| Condition Name | Chemical Name | Target Kinases (gene symbol) | Target Information Source |
|---|---|---|---|
| AKT[1] | Akt inhibitor VIII | AKT1 AKT2 AKT3 | `https://www.selleckchem.com/products/akti-1-2.html` |
| AKT[2] | MK-2206 | AKT1 AKT2 AKT3 | `https://www.selleckchem.com/products/MK-2206.html` |
| CAMK[1] | KN-93 | CAMK2A CAMK2B CAMK2D CAMK2G | `https://www.selleckchem.com/products/kn-93.html` |
| CAMK[2] | KN-62 | CAMK2A CAMK2B CAMK2D CAMK2G CAMK1 CAMK1D CAMK1G CAMK4 | `https://www.selleckchem.com/products/kn-62.html` |
| EGFR[1] | PD-168393 | EGFR | `https://www.selleckchem.com/products/pd168393.html` |
| EGFR[2] | PD-153035 | EGFR | `https://www.selleckchem.com/products/pd153035.html` |

| | | | |
|---|---|---|---|
| ERK[1] | ERK inhibitor | MAPK1 MAPK3 | https://www.sigmaaldrich.com/catalog/product/mm/328007 |
| ERK[2] | ERK inhibitor II | MAPK1 MAPK3 | https://www.sigmaaldrich.com/catalog/product/mm/328006m |
| MEK[1] | GSK-1120212 | MAP2K1 MAP2K2 | https://www.selleckchem.com/products/gsk1120212-jtp-74057.html |
| MEK[2] | U0126 | MAP2K1 MAP2K2 | https://www.medchemexpress.com/u-0126.html |
| MTOR[1] | KU-0063794 | MTOR | https://pubchem.ncbi.nlm.nih.gov/compound/Ku-0063794#section=Biomolecular-Interactions-and-Pathways |
| MTOR[2] | Torin-1 | MTOR | https://www.cellsignal.com/products/activators-inhibitors/torin-1/14379 |
| P70S6K[1] | PF-4708671 | RPS6KB1 | https://www.selleckchem.com/products/pf-4708671.html |
| P70S6K[2] | DG2 | RPS6KB1 | https://www.sigmaaldrich.com/catalog/product/mm/559274 |
| PI3K[1] | GDC-0941 | PIK3CA PIK3CB PIK3CD PIK3CG | https://www.stemcell.com/gdc-0941.html |
| PI3K[2] | PI-103 | PIK3CA PIK3CB PIK3CD PIK3CG MTOR | https://www.stemcell.com/gdc-0941.html |
| PKC[1] | Gö-6976 | PRKCA PRKCB PRKCG PRKCD PRKCZ JAK2 FLT3 | https://www.selleckchem.com/products/go6976.html https://pubmed.ncbi.nlm.nih.gov/16956345/ |
| PKC[2] | BIM-1 | PRKCA PRKCB PRKCG | https://www.selleckchem.com/products/gf109203x.html |

| | | | |
|---|---|---|---|
| ROCK[1] | H-1152 dihydrochloride | ROCK2<br>CAMK2A<br>CAMK2B<br>CAMK2D<br>CAMK2G<br>PRKG1<br>PRKG2<br>AURKA<br>PRKACA<br>PRKACB<br>PRKACG<br>PRKCA<br>PRKCB<br>PRKCB<br>PRKCG<br>PRKCD<br>PRKCE<br>PRKCH<br>PRKCQ<br>PRKCI<br>PRKCZ | `https://www.selleckchem.com/products/h-1152-dihydrochloride.html` |
| ROCK[2] | Y-27632 | ROCK1<br>ROCK2 | `https://www.selleckchem.com/products/Y-27632.html` |
